# A conserved structural logic underlies sensor–helper NLR communication in the NRC immune receptor network

**DOI:** 10.64898/2026.05.18.725810

**Authors:** AmirAli Toghani, Maián Garro, Raoul Frijters, Sophien Kamoun, Mauricio P. Contreras

**Author notes:** Authors contributed equally (AmirAli Toghani, Maián Garro).

## Abstract

NLR immune receptor networks consist of expanded disease resistance proteins (sensor NLRs) that signal via core executors of immunity known as helper NLRs. Although some sensor NLRs are thought to activate their cognate helpers via an activation-and-release mechanism, the structural basis of sensor-helper communication remains poorly understood. Here, we identify and validate sensor-helper NLR interfaces that are critical for immune activation in the NRC network of coiled-coil NLR immune receptors. Using AlphaFold 3 we predicted a high confidence model between the virus resistance protein Rx and its helper NLR NRC2. We validated the interfaces by loss and gain-of-function mutagenesis, including reconstituting a critical salt bridge through reciprocal mutations. We showed that these interfaces are conserved across the NRC network of asterid plants despite over 120 million years of divergence and validated the sensor-NRC interfaces within the common lettuce network. Structure-guided bioengineering of a lettuce sensor NLR enabled expansion of its NRC helper compatibility profile. These results are consistent with the activation-and-release model and point to bioengineering sensor-helper specificity in economically important crop species.

## Introduction

Nucleotide-binding and leucine-rich repeat (NLR) intracellular immune receptors are central components of innate immunity across plants, animals, and bacteria (*1–3*). In plants, NLRs detect pathogen-secreted effector proteins delivered into host cells to suppress immunity or reprogram host physiology, and their activation initiates robust defense responses that often culminate in programmed cell death (*1–4*). A unifying feature of NLR activation across kingdoms is their assembly into higher-order oligomeric signaling complexes, such as plant resistosomes or mammalian and bacterial inflammasomes, which transduce immune signals through diverse downstream mechanisms (*3–5*). NLR proteins can also have diverse and specialized functions. Some NLRs can function as single units, termed “singletons,” with one NLR protein mediating both ligand/effector perception and subsequent downstream signaling and cell death execution (*6*). However, NLRs can also function as receptor pairs or in higher-order configurations termed NLR networks (*3, 7*). In these cases, the paired NLRs have subfunctionalized: One NLR acts as a pathogen sensor that activates another NLR, known as helper, which mediates immune activation and disease resistance (*3, 6, 8*). Although substantial progress has been made in recent years in understanding the biochemical mechanisms of NLR activation, our understanding of the activation dynamics of paired and networked NLRs remains fragmentary. In particular, the structural basis of helper activation by sensor NLRs is still unknown (*9*).

NLRs typically exhibit a tripartite modular architecture consisting of an N-terminal signaling domain, a central nucleotide-binding and oligomerization domain, and a C-terminal leucine-rich repeat (LRR) domain (*4, 5, 10*). In plants, the N-terminal domain is most commonly a coiled-coil (CC) or Toll/interleukin-1 receptor (TIR) domain. The central domain is termed NB-ARC (NB adaptor shared by APAF-1, plant R proteins, and CED-4) and is a defining feature of this protein family. The NB-ARC domain comprises the nucleotide-binding (NB), helical domain 1 (HD1), and winged helix domain (WHD), and functions as a molecular switch that controls NLR activation (*4, 5, 10*). Structural studies of the Arabidopsis singleton NLR AtZAR1 and the *Nicotiana benthamiana* helper NbNRC2a revealed that NLR activation is accompanied by major conformational rearrangements within the NB-ARC, most notably involving rotation of the NB–HD1 module relative to the WHD (*9, 11, 12*). These rearrangements expose interfaces required for oligomerization, driving assembly into pentameric or hexameric resistosomes, respectively. Whereas resting state ZAR1 occurs as a monomer before activation, structural studies of resting state NbNRC2a and SlNRC2 revealed that these helpers exist as autoinhibited homodimers (*11, 13*). These homodimers can assemble into tetramers and even into higher-order filamentous resting-state assemblies (*13, 14*). However, how sensors engage with resting state helpers to mediate activation-associated rearrangements in paired and networked NLR systems remains unresolved.

The NRC (NLR required for cell death) network is an extensively studied example of a plant immune receptor network consisting of sensor and helper CC-NLR proteins. Found throughout Asterid species, the NRC network emerged approximately 125 million years ago and subsequently diversified into lineage-specific repertoires of NRC helper NLRs and NRC- dependent sensor NLRs (NRC-S) (*7, 15, 16*). In Solanales, for example, this diversification resulted in expanded networks comprising hundreds of NRC-S and dozens of NRC helpers, whereas other lineages such as Asterales retain more compact NRC networks with fewer components (*15–17*). Despite lineage-specific expansions, NRC0 is the only helper NLR that is broadly conserved across asterids (*15, 17*). In many species, NRC-S and their matching NRC0 are often clustered in the genome (*15, 17*).

In the NRC network, activation occurs via an activation-and-release mechanism: activated NRC-S trigger NRC helper oligomerization into resistosomes without stable incorporation of the sensor into the final helper-only wheel-like complex (*18*). Previous work demonstrated that the NB domain of NRC-S NLRs is sufficient to activate NRC helpers and can recapitulate the helper specificity of full-length sensors, implicating the NRC-S NB domain as a key activation signal of NRC helpers (*19*). The prevailing working model therefore centers on transient sensor–helper interactions as drivers of activation in the NRC network.

Despite extensive genetic and biochemical characterization, whether direct sensor-helper interactions mediate NRC helper activation and the precise interfaces that mediate this interaction remain unknown, likely due to the transient nature of these proposed activation complexes and intermediates. Decoding the molecular determinants of sensor-helper specificity in the Solanales NRC network has also been hindered by the large degree of diversification between NRC-S and NRC helpers (*15–17*). We recently reported that, in contrast to the highly expanded NRC repertoires of Solanales species, common lettuce encodes a smaller NRC network consisting of two NRC helpers and about a dozen of NRC-S with phylogenetically defined helper compatibility profiles (*16*). In particular, the close evolutionary relatedness of lettuce NRC-S coupled with their traceable helper specificities offer an opportunity to dissect how sensor-helper specificity can be encoded in this network.

Recent advances in protein structure prediction have enabled new approaches to study immune NLR biology that were previously hindered by the need for experimental structures (*20–22*). AlphaFold has allowed for the prediction of NLR-effector interaction interfaces, enabling structure-guided engineering of novel effector recognition specificities (*23–25*). Recently, AlphaFold 3 has been shown to accurately predict activated NLR resistosome assemblies, including large oligomeric immune complexes (*9, 26, 27*). Importantly, AlphaFold 3 predictions have been used to supplement cryo-EM structures of activated NLRs, as these predictions can be used to model regions that are difficult to determine experimentally, such as the membrane-associated N-terminal funnels of CC and CC_G10_ NLR resistosomes (*9, 26, 27*). These developments raise the possibility that AlphaFold 3 could be used to study additional aspects of NLR activation, such as any sensor-helper complexes that have eluded structural characterization by conventional approaches. If experimentally validated, such predictions could offer a powerful route to dissecting the structural basis of activation in the NRC network and yield insights into the molecular determinants of sensor-helper communication and compatibility.

Here, we combine AlphaFold 3-based structure prediction with extensive functional validation to define the structural basis of sensor-mediated helper activation in the NRC immune receptor network. We predicted and experimentally validated a complex between the NRC-S NLR Rx and its helper NbNRC2a, revealing a sensor–helper interface mediated by the sensor NB domain and the helper NB and LRR domains. By integrating structure-guided mutagenesis, functional assays, and comparative analyses across Asterid species, we propose that this interface is required for helper activation by sensors while also encoding sensor-helper compatibility, following a conserved structural logic that has been retained in NRC networks across Asterids. Together, these findings support a structurally informed working model in which sensor activation exposes the NB domain as an endogenous danger signal that primes helper activation via direct sensor-helper interaction, providing structural granularity to the activation-and-release mechanistic framework for signaling in the NRC networks and opening new avenues for bioengineering disease resistance by expanding sensor-helper compatibility.

## Results

### Co-immunoprecipitation fails to provide conclusive evidence for specific sensor–helper NLR associations

Since the initial characterization of the NRC network nearly 10 years ago (*7*), we have made repeated attempts to biochemically capture sensor-helper NLR interactions by co-immunoprecipitation (Co-IP) without obtaining conclusive results (*18, 19, 28*). As an illustrative example, Co-IP experiments between the NRC-S Rx and its helper NLRs NbNRC2 and NbNRC4 revealed interaction signals comparable to those obtained with the negative control NbZAR1 (unrelated to the NRC network), suggesting that these NLRs associate non-specifically under standard Co-IP conditions (**Fig. 1A**). Although these results do not rule out the possibility that Rx and its NRC helpers associate in planta, the comparable signal obtained with the negative control prevents any firm conclusions from being drawn. Together, these results underscore the limitations of Co-IP approaches for capturing transient or low-affinity sensor–helper interactions and highlight the need for alternative strategies to study the structural basis of sensor–helper communication.

**Figure 1.**
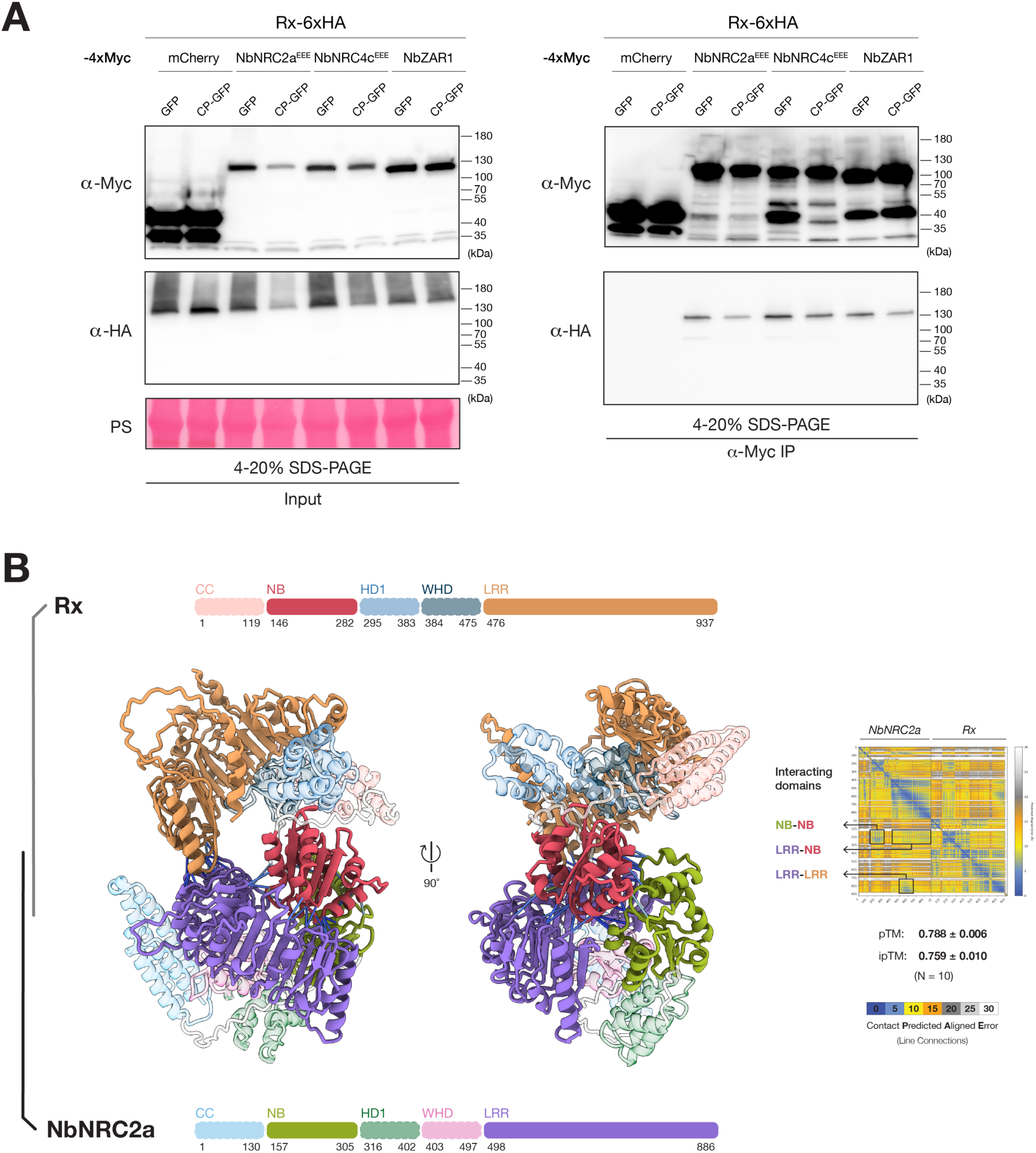
AlphaFold 3 predicts a high confidence complex between the sensor Rx and its helper NbNRC2a. (**A**) Co-immunoprecipitation assays to test the association of Rx with NbNRC2^EEE^, NbNRC4 ^EEE^, and NbZAR1 in *N. benthamiana nrc2/3/4* CRISPR mutant lines. C-terminally 6xHA-tagged Rx was co-expressed with C-terminally 4xMyc-tagged NbNRC2a ^EEE^, NbNRC4c ^EEE^, or NbZAR1 in the presence of GFP (inactive state) or GFP-tagged PVX coat protein CP (active state). Association between Rx and C-terminally 4xMyc-tagged mCherry was included as a negative control. Immunoprecipitations were performed with agarose beads conjugated to Myc antibodies (Myc IP). Total protein extracts were immunoblotted with the antisera labelled on the left. Approximate molecular weights (in kDa) of the proteins are shown on the right. Protein loading control was carried out using Ponceau stain (PS). The experiment was performed three times with similar results. (**B**) Selected views of the predicted model of Rx in complex with NbNRC2a. The predictions were run in 10 replicates with seeds 1 to 10. The model with the seed = 1 was selected as the representative model. Structures of Rx and NbNRC2a are colored based on the schematic representation of the domain architectures of each protein shown at the top and bottom, respectively. PAE plot for the representative model as well as the average and SD values for ipTM and pTM (N = 10) are shown on the right.

### AlphaFold 3 predicts a high-confidence model for the NRC-S Rx in complex with its helper NbNRC2a

In the activation-and-release model, NRC-S activate downstream NRC helper NLRs without stably incorporating into the final helper homooligomeric resistosome (*9, 18, 19, 27–29*). This implies that sensor–helper communication may occur via direct but transient interactions that are difficult to capture experimentally. To gain structural insight into this putative activation intermediate, we used AlphaFold 3 to predict the structure of the NRC-S Rx in complex with its NRC helper NbNRC2a (**Fig. 1**). We modelled a heterodimeric Rx–NbNRC2a complex using 10 independent AlphaFold 3 predictions with seeds 1 through 10, all of which converged on a similar interaction mode. The resulting models exhibited high confidence metrics, including consistently high predicted local distance difference test (pLDDT) values, a mean predicted template modeling score (pTM) of 0.788, a mean interface pTM (ipTM) of 0.759 (**Fig. 1B, Fig. S1**).

### Rx-NbNRC2a complex is mediated through NB-NB, NB-LRR, and LRR-LRR interactions

All 10 AlphaFold 3 models reproducibly positioned Rx and NbNRC2a in the same relative orientation, revealing a highly consistent interaction topology between the two proteins (**Fig. 2A-B**, **Fig. S1**). In these models, the NB domain of the NRC-S Rx engages both the NB and LRR domains of the helper NbNRC2a, forming two major interaction surfaces that we define as the NB–NB and NB–LRR interfaces. Together, these interfaces comprise a total of 185 inter-residue contacts and bury a combined surface area of 2026.4 Å² (**Data S1**).

**Figure 2.**
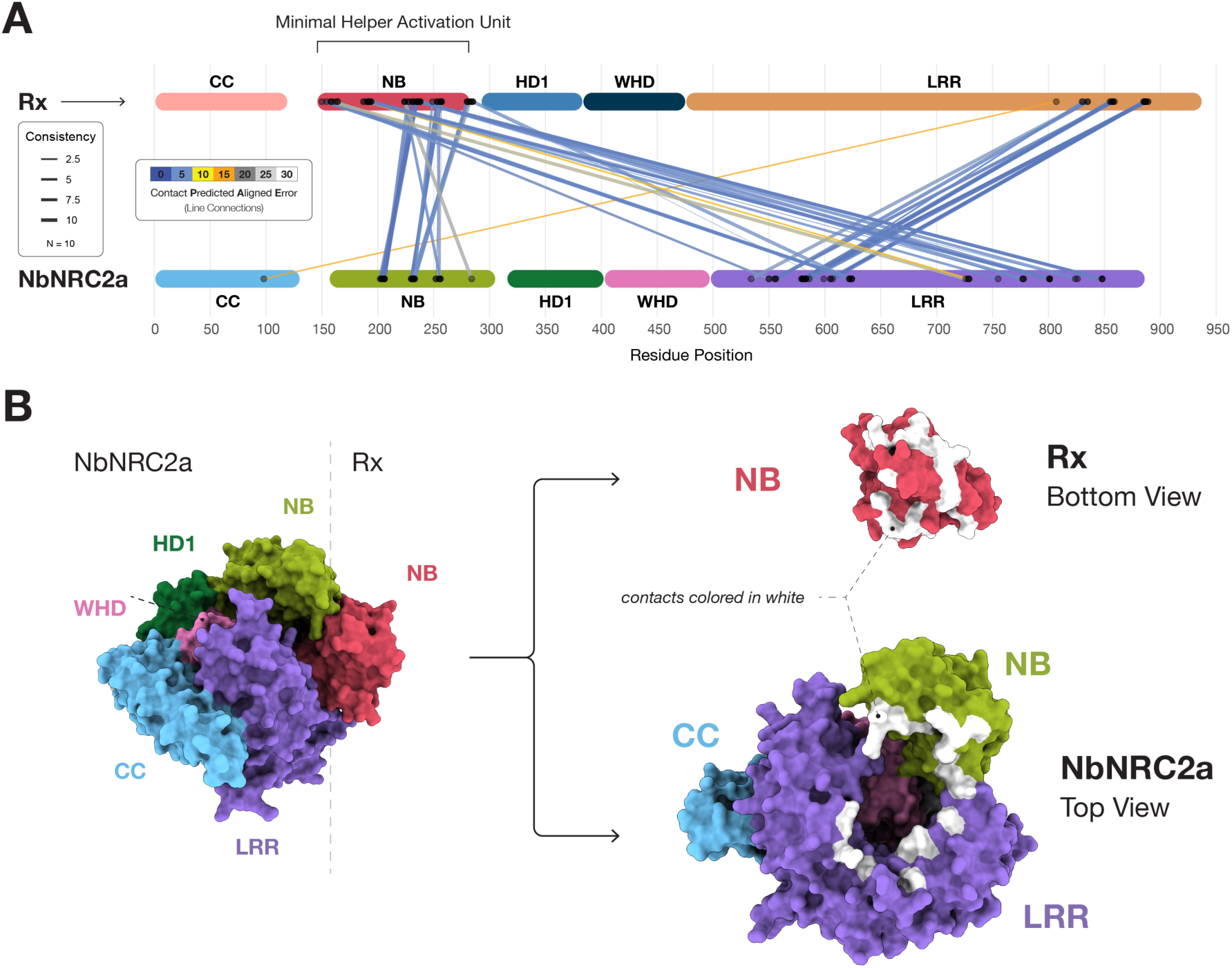
The predicted Rx-NbNRC2a complex is formed by NB-NB, NB-LRR, and LRR-LRR interactions. (**A**) Schematic representation of the domain architectures of Rx and NbNRC2a. Lines indicate contacts between the two proteins at the connected positions, as predicted by AlphaFold 3. The color of the lines indicates the contact Predicted Aligned Error for that interaction and the line thickness indicates the consistency of that contact across all 10 modeling replicates. The sensor NB domain, which is the minimal unit required for NRC helper activation, is highlighted in brackets. (**B**) Open book representation of the isolated Rx NB domain (Rx^NB^) in complex with NbNRC2a. The structures are colored based on the schematic representation of the domain architectures shown in panel (**A**). Surfaces involved in mediating the predicted interaction are shown in white.

Notably, the predicted involvement of the Rx NB domain in both interfaces is consistent with previous functional data showing that the NB domain of NRC-S is the minimal unit required to activate NRC helper oligomerization and cell death (**Fig. 2A**) (*19*). In addition to these major interfaces, the models revealed a third interaction interface mediated by the C-terminal region of the Rx LRR domain contacting the LRR domain of NbNRC2a, which we define as the LRR–LRR interface. This interface is comparatively smaller in size, with 50 contacts and a buried surface area of 638.41 Å² (**Data S1**).

Because the NB domain alone is sufficient to activate NRC helpers (*19*), we next modeled the isolated NB domain of Rx (Rx^NB^) in complex with NbNRC2a using AlphaFold 3 (**Fig. 2B**). These predictions also yielded high-confidence models in which Rx^NB^ engages NbNRC2a through the same interface observed in the full-length Rx predictions. Superposition of the Rx^NB^–NbNRC2a and full-length Rx–NbNRC2a models showed close agreement at the predicted sensor–helper interface (Unpruned RMSD = 0.896 Å; **Fig. S2**), indicating that the predicted NB-mediated interaction is independent of the remainder of the sensor protein.

### Structure-guided mutations in the predicted sensor-helper interaction interface abolish cell death triggered upon Rx^NB^ activation of NbNRC2a

To test the degree to which the AlphaFold 3-predicted Rx–NbNRC2a interface is functionally relevant, we designed a series of structure-guided loss-of-function mutations within the NB domain of Rx. Based on the predicted interaction surface, we subdivided the Rx NB domain into five discrete stretches of amino acids (Stretches 1–5; **Fig. 3A**). Stretches 1, 2, and 5 contribute primarily to the NB–LRR interface, stretch 4 maps to the NB–NB interface, and stretch 3 participates in both interfaces (**Fig. 3A-B**).

**Figure 3.**
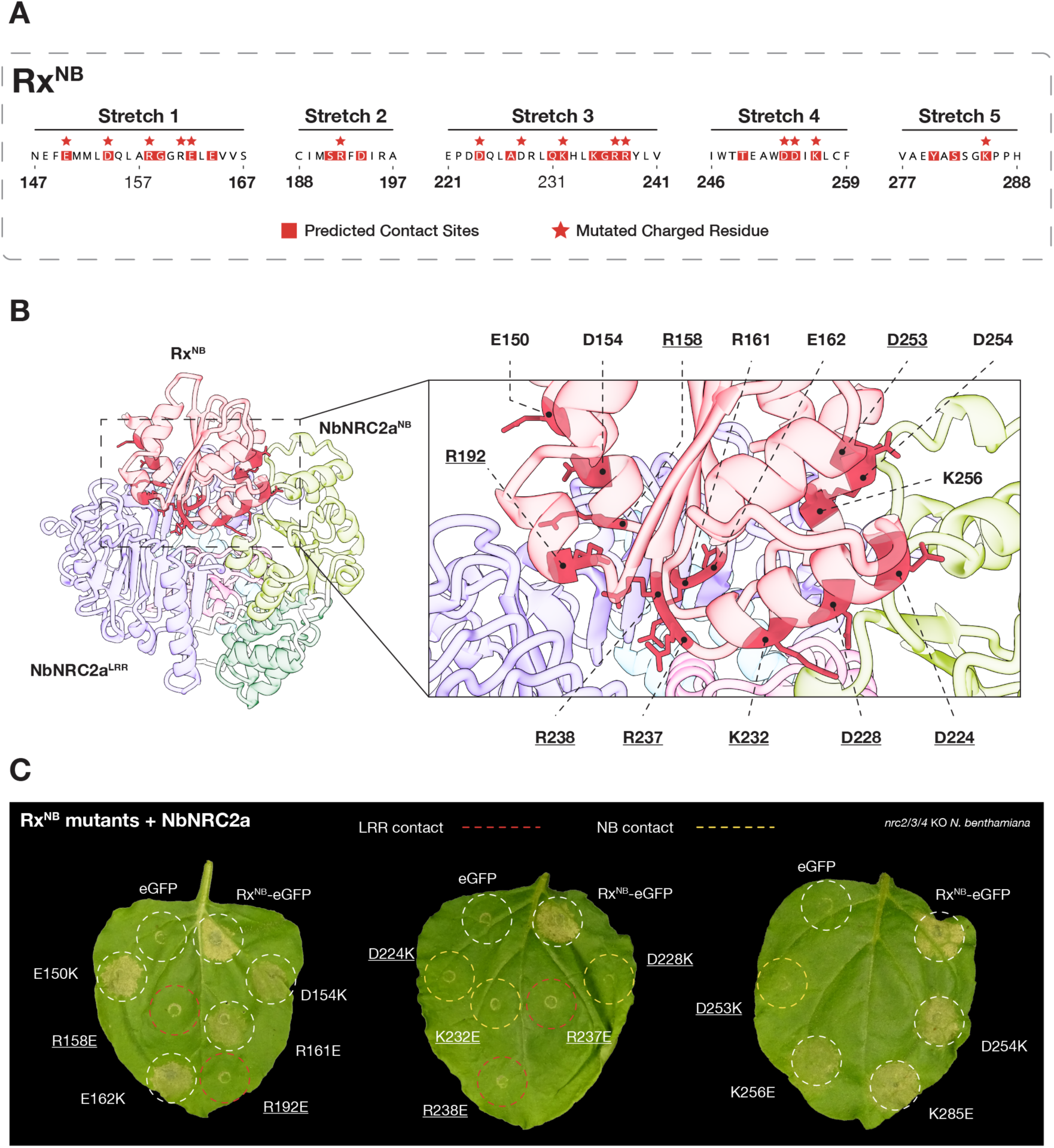
Structure-guided Rx NB domain point mutants in the predicted sensor-helper interface are unable to activate NbNRC2a. (**A**) Amino acid sequence of the 5 stretches within the NB domain or Rx that are involved in the Rx-NbNRC2a interface, specifically in the NB-NB and NB-LRR interfaces. Predicted contact sites are indicated with red shading. Charged residues that were selected for mutation to the opposite charge are indicated with red stars. (**B**) Close-up of the Rx^NB^-NbNRC2a interface, with all the mutated Rx residues highlighted. Residues that led to loss of NbNRC2a activation are underlined. Structures are colored according to the schematic representations shown in Fig. 1 and Fig. 2A. (**C**) Photo of representative leaves from *N. benthamiana nrc2/3/4* CRISPR mutant lines showing cell death after co-expression of NbNRC2a with different Rx^NB^-eGFP variants generated. Wild-type Rx^NB^-eGFP and free eGFP were co-expressed with NbNRC2a as positive and negative controls for cell death, respectively. For mutations that led to loss of NbNRC2a activation, color of the dotted outline indicates if that amino acid in Rx was predicted to make contacts with the NbNRC2a NB (Yellow) or LRR (Red). The experiment consisted of three biological replicates. See **Fig. S3** for quantitative analysis of the cell death phenotypes and for WB analysis of all variants tested.

Within these five stretches, we identified 15 charged residues (arginine or lysine, and aspartate or glutamate) that were consistently predicted to form inter-residue contacts with NbNRC2a across independent AlphaFold 3 replicates (**Fig. 3A, Data S1**). To disrupt the predicted interface, each residue was individually substituted with an amino acid of opposite charge. These substitutions were introduced as point mutations into Rx^NB^, which we previously showed is the minimal NRC-S unit capable of activating NRC helpers (*19*), and the resulting variants were tested for their ability to induce cell death in leaves of *nrc2/3/4 N. benthamiana* CRISPR-derived mutant lines complemented with NbNRC2a (**Fig. 3C**, **Fig. S3B**).

Eight of the 15 Rx^NB^ variants completely lost the ability to trigger NbNRC2a-dependent cell death. Four of these loss-of-function mutations (D224K, D228K, K232E, and D253K) mapped to residues within stretches 3 and 4 that mediate NB–NB interactions with NbNRC2a (**Fig. 3A-B**). The remaining four loss-of-function variants (R158E, R192E, R237E, and R238E) mapped to residues within stretches 1, 2, and 3 involved in NB–LRR interactions (**Fig. 3B-C**). One additional variant in stretch 1 (R161E) displayed a partial reduction in NbNRC2a activation, whereas the remaining five variants retained cell death–inducing activity comparable to wild-type Rx^NB^ (**Fig. 3C**). Immunoblot analysis showed that all eight loss-of-function Rx^NB^ variants accumulated to levels comparable to the wild-type protein, indicating that their impaired ability to activate NbNRC2a is not due to reduced protein stability (**Fig. S3**). Together, these results demonstrate that both NB–NB and NB–LRR predicted interaction interfaces are required for NbNRC2a activation by Rx.

### Four Rx NB domain residues at the predicted sensor-helper interface are required for activation of NbNRC2a, NbNRC3 and NbNRC4c

In leaves of wild-type *N. benthamiana*, the NRC-S Rx can initiate programmed cell death via multiple helper NLRs, including NbNRC2a, NbNRC3, and NbNRC4c (*30*). To determine the degree to which the Rx–NbNRC2a interface identified above is also important for Rx-mediated activation of additional NRC helpers, we used AlphaFold 3 to model Rx in complex with NbNRC3 and NbNRC4. Across 10 independent predictions for each complex, AlphaFold 3 consistently placed Rx and each helper in a similar relative orientation, yielding predicted interactions through NB-NB and NB-LRR interfaces like those observed in the Rx–NbNRC2a models (**Fig. S4**). Moreover, to rule out non-specific prediction of confident complexes by AlphaFold 3 for Rx and NRC helpers, we modelled Rx–AtZAR1 and AtZAR1–NbNRC2a complexes, with AtZAR1 serving as a negative control. Across all 10 replicates for each complex, we observed no confident contacts, and ipTM values did not exceed 0.3, indicating poor model quality (**Fig. S4**).

To directly dissect helper-specific contributions of each of the residues mutated in each Rx^NB^ variant, we expressed the panel of 15 variants in leaves of *nrc2/3/4* CRISPR mutant *N. benthamiana* lines individually complemented with NbNRC3 or NbNRC4. Four Rx^NB^ variants R192E, D228K, K232E, and R238E were unable to activate either NbNRC3 or NbNRC4. In contrast, the remaining variants exhibited differential effects: R158E retained near–wild-type activation of NbNRC3 but showed reduced signaling via NbNRC4; D253K activated NbNRC3 but not NbNRC4: all other variants, including D224K and R237E, retained the ability to signal through both helpers (**Fig. S5**).

To further confirm these results, we tested all variants that exhibited a loss-of-function phenotype with at least one helper (R158E, R192E, D224K, D228K, K232E, R237E, R238E and D253K) and tested these in WT *N. benthamiana*, which expresses all three NRC helpers endogenously (**Fig. S5**). Of the eight variants tested, four variants (R192E, D228K, K232E and R238E) completely lost the ability to induce cell death in wild-type plants, consistent with the observation that these mutants were impaired in their ability to signal with all helpers tested (**Fig. S5**). The remaining loss-of-function variants displayed partial or no impairment in wild-type *N. benthamiana*, consistent with their differential contributions to helper activation depending on which NRC is engaged. Together, these results define two functional classes of residues in the helper-interacting surface of the Rx NB domain. A core set of residues is required for activation of all NRC helpers tested, whereas additional residues contribute to signaling in a helper-specific manner.

### Two residues in the NB domain of NbNRC2 are required for activation by Rx^NB^

To further investigate the functional relevance of the predicted Rx–NbNRC2a interaction interface, we generated structure-guided mutations in NbNRC2a targeting residues within the NB–NB and NB–LRR interfaces (**Fig. 4A**). Using the same strategy applied to Rx, we selected nine charged residues in NbNRC2a that were consistently predicted by AlphaFold 3 to form contacts with the NB domain of Rx across independent modeling replicates. Each residue was individually substituted with an amino acid of opposite charge to disrupt the predicted interaction surface.

**Figure 4.**
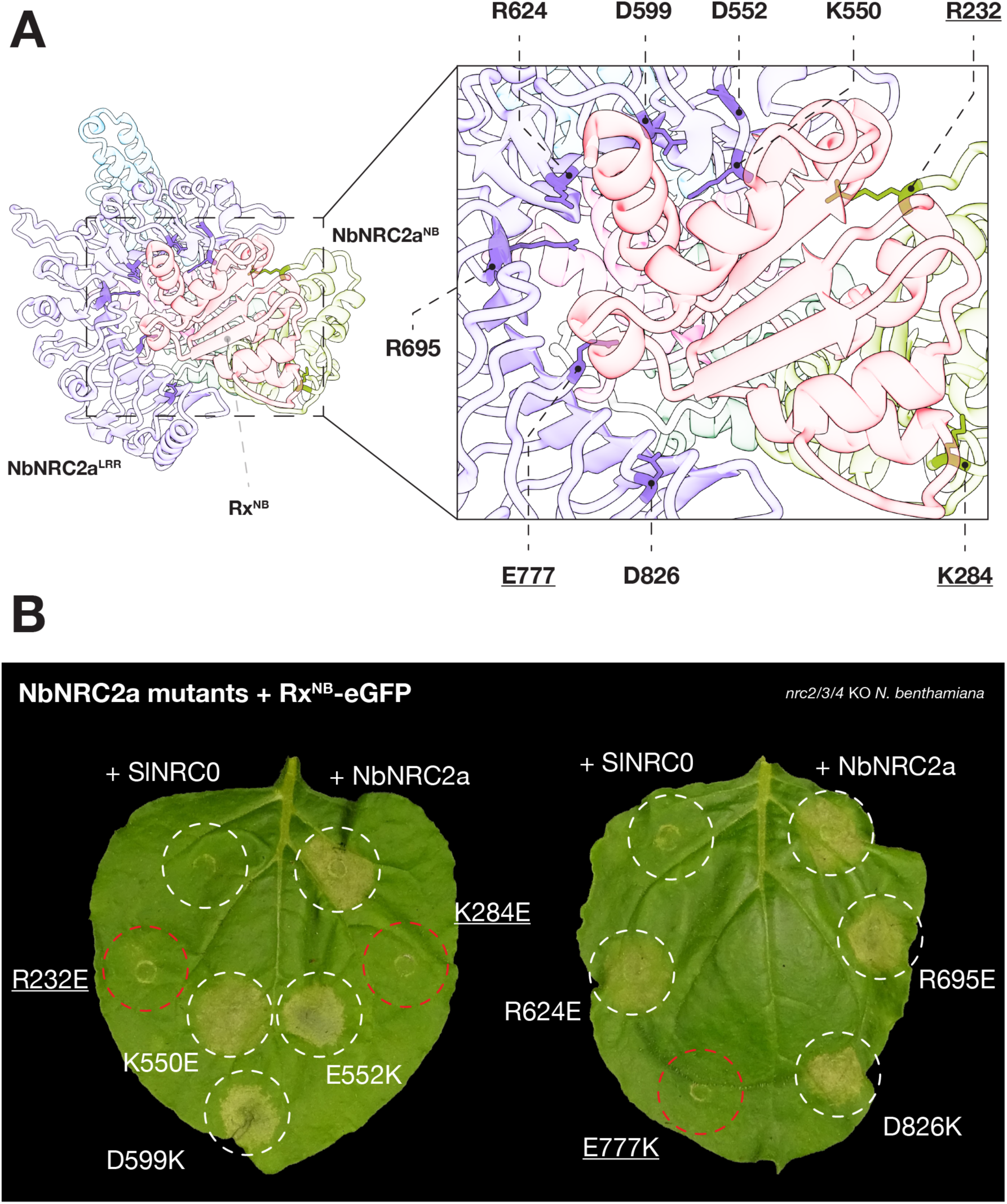
Structure-guided NbNRC2a interface mutations impair Rx^NB^-dependent activation. (**A**) Selected view of the AlphaFold 3-predicted Rx–NbNRC2a interface highlighting NbNRC2a residues targeted for charge-swap mutagenesis (NB–NB and NB–LRR contact sites). (**B**) Representative *nrc2/3/4* CRISPR mutant *N. benthamiana* leaf images showing cell death after co-expression of Rx^NB^-eGFP with NbNRC2a (WT) or the indicated NbNRC2a point mutants. NbNRC2a (WT) and SlNRC0 were included as positive and negative controls for Rx^NB^- dependent cell death, respectively. The experiment was repeated three times with at least six technical replicates per repeat, with similar results. See **Fig. S6** for quantitative analysis of the cell death phenotypes and for WB analysis of all variants tested.

The nine NbNRC2a variants were individually tested for their ability to complement Rx^NB^–induced cell death in leaves of *nrc2/3/4 N. benthamiana* CRISPR lines (**Fig. 4B**, **Fig. S6B**). Three of the nine variants failed to restore Rx^NB^-mediated cell death. Two of these loss-of-function mutations (R232E and K284E) mapped to residues within the NB domain of NbNRC2a predicted to participate in NB–NB interactions with Rx. The third loss-of-function mutation (E777K) mapped to the LRR domain and disrupted the predicted NB–LRR interface. The remaining six NbNRC2a variants retained the ability to support Rx^NB^-dependent cell death at levels comparable to wild-type NbNRC2a.

Immunoblot analysis revealed that the R232E and K284E NbNRC2a variants accumulated to levels comparable to wild-type NbNRC2a, indicating that their impaired ability to support cell death is not due to reduced protein accumulation (**Fig. S6**). In contrast, although the NbNRC2a E777K variant was consistently detectable, it showed reduced accumulation, which may contribute to its loss-of-function phenotype. Together, these results provide reciprocal functional support for the AlphaFold 3-predicted sensor–helper interface and indicate that interactions at this interface are important for NbNRC2a activation by upstream NRC-S such as Rx.

### Reconstituting an NB-NB salt bridge through reciprocal mutations rescues Rx-mediated NbNRC2a cell-death

Although the loss-of-function experiments described above support the functional relevance of the AlphaFold 3-predicted Rx–NbNRC2a interface, such phenotypes can in principle arise from indirect effects such as protein misfolding. To directly challenge the hypothesis that specific inter-molecular contacts within the predicted interface mediate sensor–helper activation, we focused on residues predicted to form inter-molecular salt bridges across the sensor–helper interface using AlphaFold 3 modeling. These included a salt bridge between D224 in the Rx NB domain and K284 in the NbNRC2a NB domain (**Fig. 5A**), as well as a R192 and R238 in the Rx NB domain and E777 in the NbNRC2a LRR domain (**Fig. S7A**). All five of these residues resulted in loss-of-function when mutated (**Fig. 3**, **Fig. 4**).

**Figure 5.**
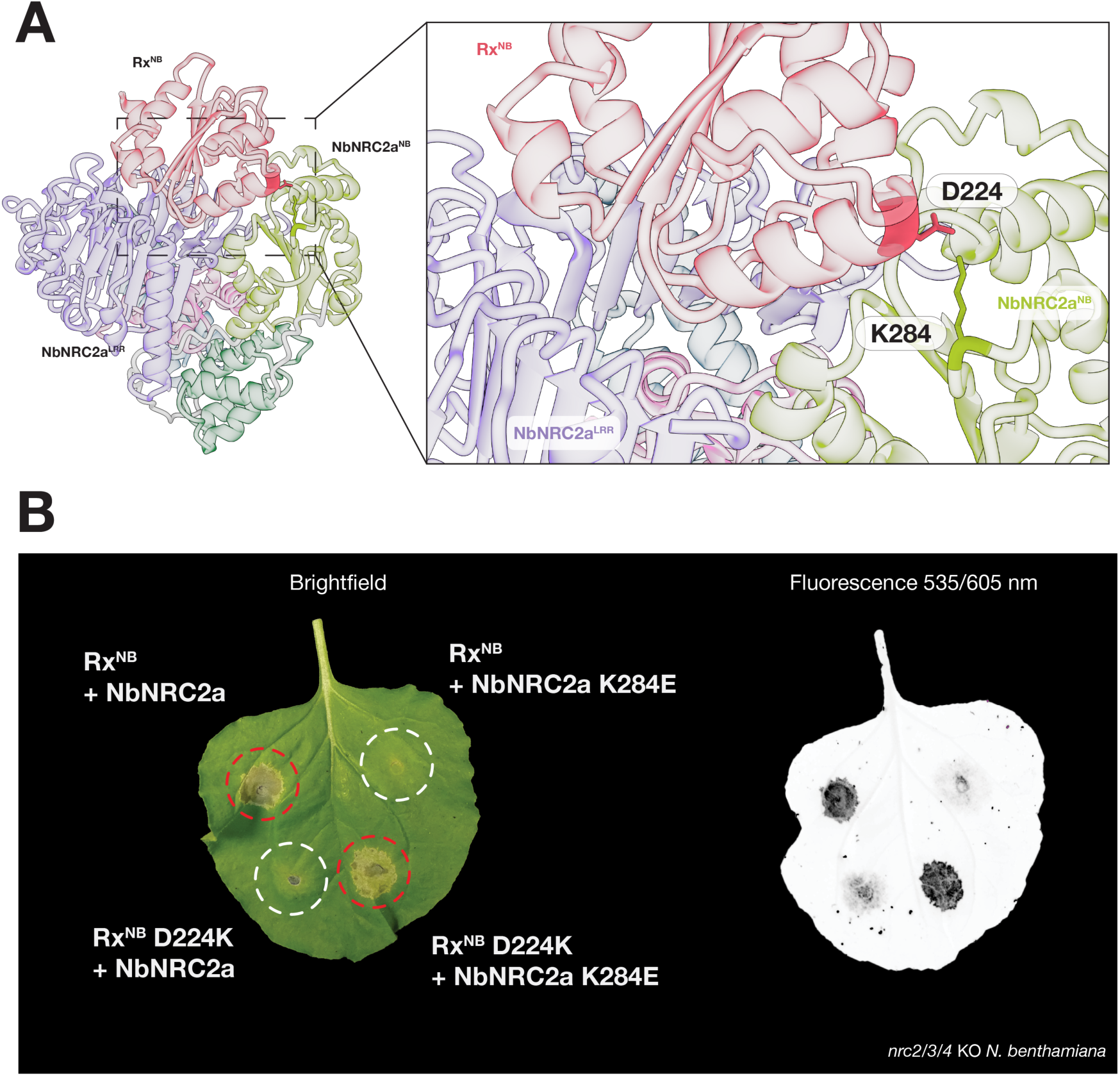
Reciprocal charge swaps restore Rx^NB^–NbNRC2a signaling. (**A**) Selected AlphaFold 3-predicted salt bridge across the Rx^NB^–NbNRC2a interface highlighting the residue pair targeted for reciprocal charge-swap mutagenesis. (**B**) Representative *nrc2/3/4* CRISPR mutant *N. benthamiana* leaf images (bright-field and UV) showing cell death following co-expression of Rx^NB^ with NbNRC2a (positive control), Rx^NB^ or NbNRC2a single charge-swap mutants paired with the corresponding WT partner, or matched Rx^NB^/NbNRC2a reciprocal charge-swap combination. The experiment was repeated three times with at least six technical replicates per repeat, with similar results.

We hypothesized that if these residue pairs form biologically relevant salt bridges, then simultaneous inversion of charge on both sides of the interface should restore the interaction and rescue NRC helper activation. To test this, we performed reciprocal charge-swap experiments by co-expressing matched Rx and NbNRC2a variants in cell death assays in leaves of *nrc2/3/4* CRISPR mutant *N. benthamiana* lines (**Fig. 5B**). In agreement with this hypothesis, co-expression of Rx^NB-D224K^ with NbNRC2a^K284E^ resulted in a strong cell death response comparable to that observed upon co-expression of wild-type Rx^NB^ and NbNRC2a. In contrast, co-expression of wild-type Rx^NB^ with NbNRC2a^K284E^ or Rx^NB-D224K^ with wild-type NbNRC2a failed to trigger cell death, consistent with the loss-of-function phenotypes observed for the individual mutants (**Fig. 5B**).

We next tested the second predicted salt bridge involving the Rx NB domain and the NbNRC2a LRR domain. Co-expression of Rx^NB-R192E^ with NbNRC2a^E777K^ resulted in a partial restoration of cell death, although the response did not reach the levels observed for the wild-type pair. In contrast, co-expression of RxNB^R238E^ with NbNRC2a^E777K^ did not induce visible cell death (**Fig. S7B**). Notably, the NbNRC2a^E777K^ variant accumulates at reduced levels compared to wild-type NbNRC2a, which may limit the extent of functional rescue observed in these assays (**Fig. S6**).

Together, these results provide strong experimental support for the AlphaFold 3- predicted salt bridge between D224 in Rx and K284 in NbNRC2a, and more limited support for an additional interaction between the Rx NB domain and the NbNRC2a LRR domain. The ability to restore Rx^NB^-mediated NbNRC2a activation through reciprocal charge-swap mutations provides compelling functional evidence that specific inter-molecular contacts within the predicted sensor–helper interface are required for helper activation.

### NB-domain interactions, but not the LRR–LRR interface, are required for helper activation by full-length Rx

To what extent do the results obtained with Rx^NB^ translate to the full-length Rx protein? We investigated this using structure-guided mutations in full-length Rx and assesssed the contribution of the interaction interface involving the LRR domain of both proteins as predicted by AlphaFold 3 (**Fig. 1**, **Fig. 2**). Three full-length Rx variants were constructed: Rx_NB-int_, in which the four NB-domain residues R192, D228, K232, and R238 previously identified as required for activation of all NRC helpers were mutated simultaneously; Rx_LRR-int_, in which 4 residues (E807K, D831K, D856K and D885K) within the Rx LRR domain predicted by AlphaFold 3 to contact the NbNRC2a LRR domain were mutated; and Rx_NB/LRR-int_, which combines both sets of mutations in cis. These Rx variants were tested for their ability to activate NbNRC2a following co-expression of the PVX CP.

Both Rx_NB-int_ and Rx_NB/LRR-int_ failed to trigger NbNRC2a-dependent cell death upon PVX CP recognition. In contrast, Rx_LRR-int_ retained the ability to respond to PVX CP and induce NbNRC2a-dependent cell death (**Fig. S8**). These results indicate that NB–NB and NB–LRR interactions are required for helper activation in the context of both the minimal Rx^NB^ system and full-length Rx, whereas the predicted LRR–LRR sensor-helper interface is dispensable for helper activation.

### Rx models in resting and NbNRC2a-bound conformations mirror structural transitions observed during helper activation

We noted that the AlphaFold 3 structural models of Rx in its resting and NbNRC2a-bound forms are different. This prompted us to perform further comparative structural analyses to investigate the extent to which these predictions reflect conformational changes associated with Rx activation. We predicted resting-state Rx with AlphaFold 3 and obtained high confidence models across 10 independent replicates (**Fig. 6A**). In these models, Rx adopts an overall conformation similar to NbNRC2a protomers within the resting-state homodimeric complex (*11*). This includes similar autoinhibitory interactions involving the NB and adjacent domains (**Fig. 6, Fig. S9**; RMSD = 1.235 Å between resting state Rx and NbNRC2a). In contrast, Rx predicted in complex with NbNRC2a adopts a markedly different conformation. The NbNRC2a-bound Rx model exhibits an overall domain arrangement closely resembling that of an activated NbNRC2a protomer within the hexameric resistosome (*9*) (**Fig. 6; Fig. S9**; RMSD = 1.076 Å between activated Rx and NbNRC2a).

**Figure 6:**
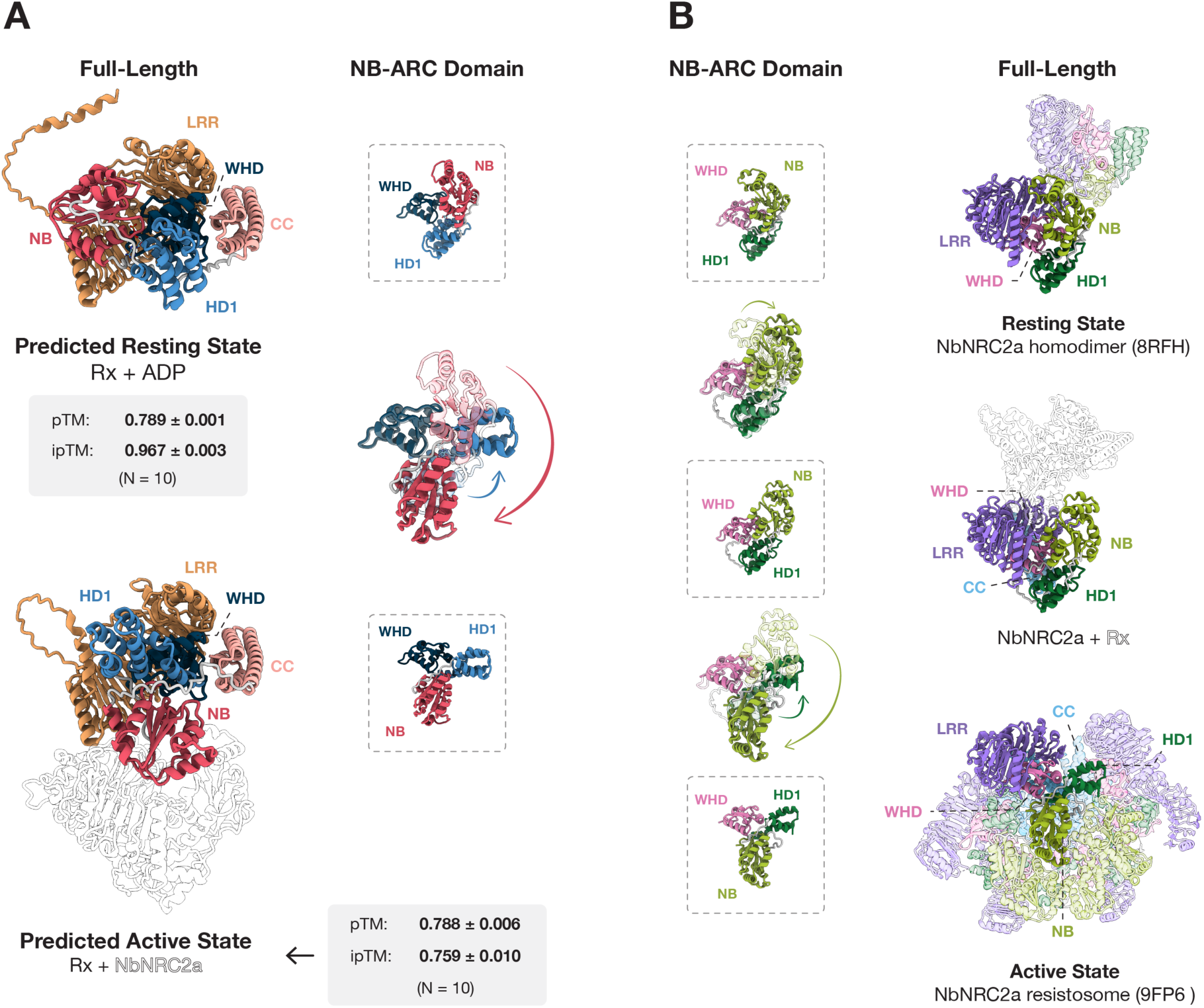
AlphaFold 3-predicted Rx conformations mirror the structural transitions observed during NbNRC2a activation. (**A**) Comparison of AlphaFold 3-predicted full-length Rx structures in the resting state (Rx + ADP, top) and sensor- helper complex state (Rx + NbNRC2a, bottom). Both predictions were run with 10 independent seeds; the seed = 1 model is shown as a representative. Mean pTM and ipTM values (± SD, N = 10) are indicated for each prediction. Protein structures are colored by domain as in Fig. 1. For the active state prediction, NbNRC2a is shown in white. The central panels show isolated NB-ARC domain views of each conformation, with arrows indicating the direction of NB–HD1 module rotation relative to the WHD between the resting and complex-bound states. (**B**) Comparison of NbNRC2a NB-ARC domain (center) and full-length structures (right) across three conformational states: the experimentally determined resting-state homodimer (PDB: 8RFH, top), the AlphaFold 3-predicted NbNRC2a conformation when bound to Rx (middle), and the experimentally determined active-state hexameric resistosome (PDB: 9FP6, bottom). The predicted Rx-bound NbNRC2a conformation is colored by domains as in Fig. 1. Arrows indicate the progressive rotation of the NB–HD1 module relative to the WHD across states, illustrating that sensor binding is predicted to prime the helper toward an intermediate conformation preceding full activation and resistosome assembly.

Comparison of the predicted resting and NbNRC2a-bound Rx structures reveals pronounced rearrangements within the NB-ARC, including a rotation of the NB–HD1 module relative to the WHD that mirrors the conformational change observed during NbNRC2a activation (**Fig. 6A-B**). Together, these observations suggest that Rx activation may involve structural transitions analogous to those undergone by NRC helper NLRs, consistent with their proposed shared evolutionary origins.

### NRC-S binding may prime NRC helper activation through NB-domain displacement and homodimer destabilization

We probed the conformation of NbNRC2a when predicted in complex with Rx. In these AlphaFold 3 models, NbNRC2a adopts a conformation distinct from the resting state, with the NB domain partially rotated outward relative to its position in the resting state NbNRC2a homodimer structure (**Fig. 6B**). This configuration is reminiscent of a “primed” activation intermediate previously described for the NLR AtZAR1, in which partial outward rotation of the NB domain precedes full activation and oligomerization (PDB 6J5V; **Fig. S10**). These observations suggest a model in which binding of activated Rx to NbNRC2a via NB–NB and NB–LRR interactions promotes a primed helper conformation by displacing the NbNRC2a NB domain. Such displacement could facilitate subsequent conformational changes within the NB-ARC, potentially enabling nucleotide exchange and the extensive NB–HD1 rotation required for NbNRC2a oligomerization and resistosome assembly (**Fig. 6B**).

Consistent with a model in which NRC-S binding engages a single helper protomer, superposition of the predicted Rx–NbNRC2a complex onto the experimentally determined NbNRC2a resting-state homodimer (PDB: 8RFH) reveals extensive steric clashes between Rx and the second, Rx unbound, NbNRC2a protomer (**Fig. S11**). These clashes indicate that the predicted sensor–helper binding mode is geometrically incompatible with NbNRC2a homodimerization, suggesting that NRC-S binding to NRC helpers may trigger homodimer dissociation. Together with the NB-domain displacement described above, these findings point to a mechanism in which NRC-S engagement may simultaneously destabilize the resting-state NRC helper homodimer and prime the bound protomer for activation. While these conclusions are based on structural predictions, they are consistent with the experimentally validated sensor– helper interface described above and provide a mechanistic framework for how effector- activated NRC-S could directly initiate NRC helper activation.

### NRC sensor-helper pairs are predicted to engage through a conserved structural logic across Asterids

Having established experimental support for the predicted Rx–NbNRC2a complex in Solanales, we next asked whether sensor–helper communication in the NRC network relies on a conserved interaction geometry across Asterid diversity. We used AlphaFold 3 to predict sensor–helper complexes from representative species spanning four additional asterid orders beyond Solanales: Garryales (*Eucommia ulmoides*), Dipsacales (*Sambucus nigra*; elderberry), Ericales (*Actinidia deliciosa*; kiwifruit), and Asterales (*Lactuca sativa*; lettuce). We also included Caryophyllales (*Beta vulgaris*; sugar beet) as the only non-asterid order known to contain NRC sequences (**Fig. 7**).

**Figure 7:**
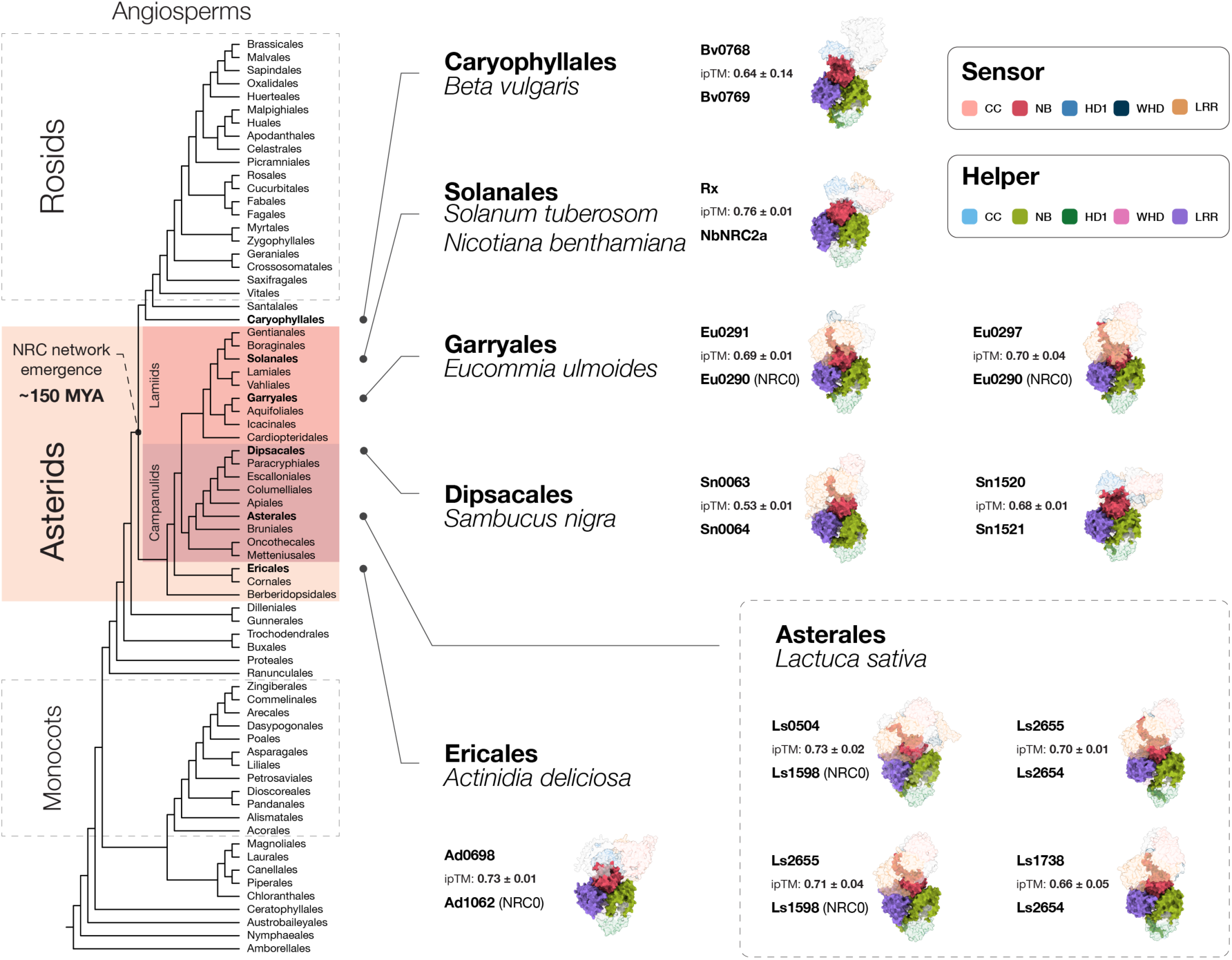
Sensor-helper interfaces are conserved in the NRC network across all Asterids. Phylogenetic tree of angiosperms obtained from Zuntini et al. (2024) (*32*). The NRC networks has emerged approximately 150 million years ago (MYA). Asterid orders in addition to Caryophyllales for which sensor–helper complexes were modeled are indicated in bold. Representatives from Caryophyllales (*Beta vulgaris*), Garryales (*Eucommia ulmoides*), Dipsacales (*Sambucus nigra*), Asterales (*Lactuca sativa*), Ericales (*Actinidia deliciosa*), are connected by lines to their corresponding AlphaFold 3-predicted structures. For each pair, sensor and helper domains colored based on domain boundaries. Mean ipTM values (± SD, N = 10) is shown for each predicted complex.

Through phylogenetic analysis and based on prior knowledge of NRC0 and NRC6 helpers and their associated NRC-S, we selected NRC sensor–helper pairs located in close genomic proximity in each species, as these are likely to be functionally linked (*15, 31*). For lettuce, we used representatives from its recently experimentally characterized NRC network (*16, 17*). In kiwifruit, where only the NRC0 helper clade is present, we assumed that the selected NRC-S signals through the sole NRC helper.

Although NRC-S and NRC helper sequences are phylogenetically diverged across these lineages, AlphaFold 3 consistently placed sensors and helpers in a similar relative orientation (**Fig. 7, Fig. S12;**). In all predicted pairs, the NB domain of the NRC-S engaged the NB and LRR domains of the helper, revealing a conserved structural logic defined by a shared binding mode and interaction topology rather than by conservation of primary sequence (**Fig. S12C, Fig. S13, Fig. S14**). Notably, while this NB-centered structural logic was observed across all predicted sensor–helper pairs, including NRC0 and its corresponding NRC-S pair, the additional LRR–LRR interface of the Rx–NbNRC2a complex was not universally present (**Fig. S13, Fig. S14**). This variability is consistent with our functional data indicating that the LRR–LRR interface is dispensable for helper activation in the Rx-NbNRC2a system (**Fig. S8**). Together, these predictions suggest that NRC network sensor–helper activation may be governed by a structural logic conserved across NRC-S and NRC sequences that involves conservation of domain orientation and binding mode. We propose that this structural logic defines a common architectural framework for sensor–helper communication across extensively diversified NRC networks.

### Structure-guided swapping of predicted sensor–helper interaction surfaces expand helper compatibility in the lettuce NRC network

To functionally test the extent to which the conserved sensor/helper structural logic is biologically relevant beyond the Rx–NbNRC2a system, we focused on the phylogenetically distant *Lactuca sativa* NRC network which is separated from the Solanales NRC network by an estimated 102 million years of evolution (*33*). Based on our structure–function analysis of Rx, we hypothesized that the NB-domain–centered interaction surfaces underlying this conserved structural logic are also required for NRC helper activation in lettuce. Furthermore, we reasoned that if these interaction surfaces also encode helper specificity, then transferring interface elements from a promiscuous to a more restricted NRC-S should expand its helper compatibility.

To test this hypothesis, we focused on four interface regions in lettuce NRC-S. These four regions are equivalent to the stretches in Rx previously identified as being involved in helper activation (Stretches 1–4; **Fig. 3A**). We selected the Clade 2 NRC-S Ls2655, which activates both lettuce helpers, as an interface donor. Chimeric sensors were generated in which individual stretches, or combinations thereof, from Ls2655 were introduced into either the Clade 1 NRC-S Ls0504 (activates LsNRC0) or the Clade 3 NRC-S Ls1738 (activates Ast-LsNRC1) (**Fig. 8A–C**). Because cognate effectors for these sensors are unknown, constitutively active variants carrying D to V substitutions in the conserved MHD motif were used to assay helper activation (Ls0504^D436V^ and Ls1741^D489V^).

**Figure 8:**
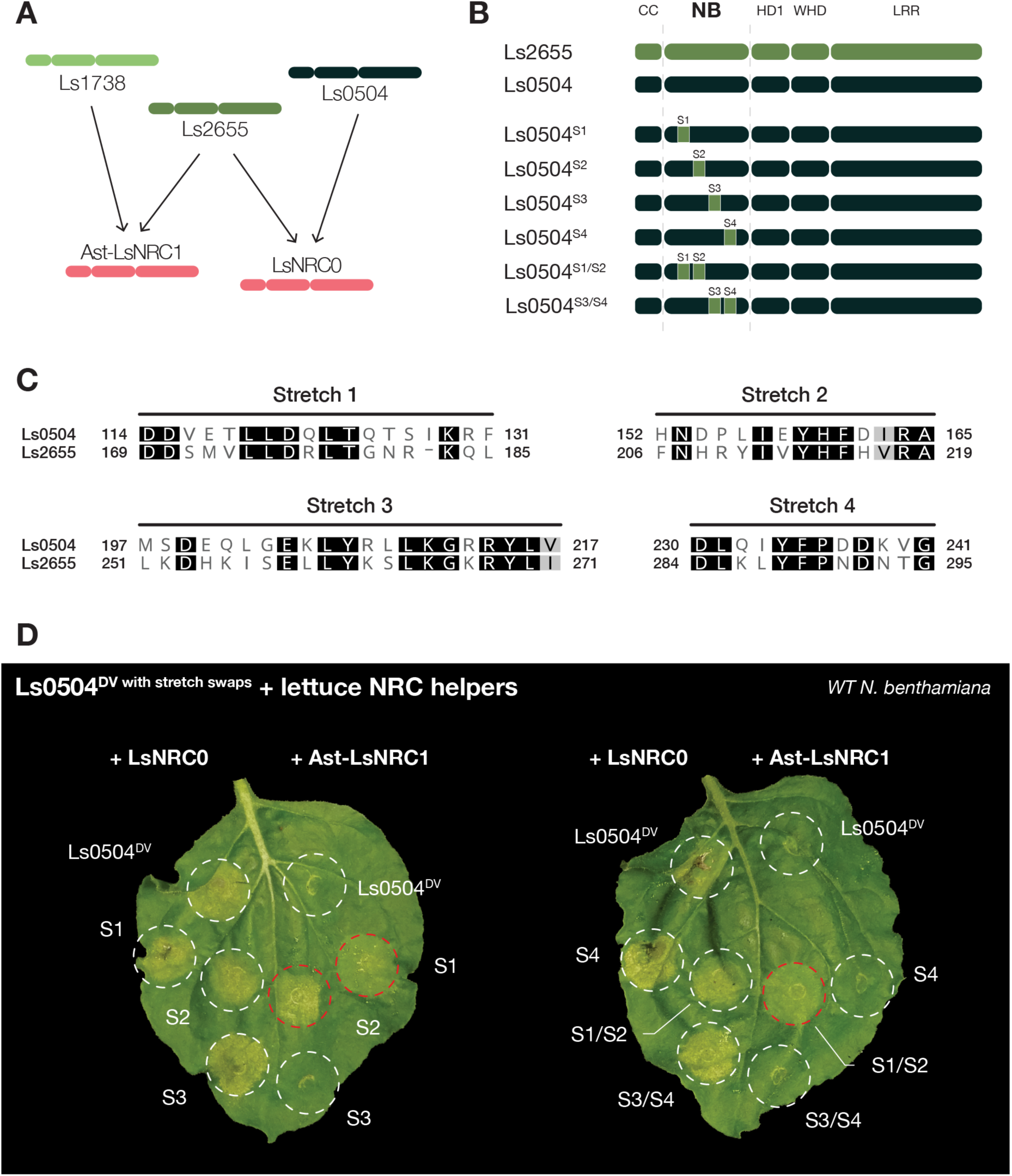
Interface swaps in Ls0504 expand helper compatibility. (**A**) Schematic of sensor–helper compatibility for representative *Lactuca sativa* NRC network NRC-S and NRC helpers used in this experiment. Arrows indicate functional signaling relationships: the Clade 1 sensor Ls0504 signals exclusively through LsNRC0, the Clade 3 sensor Ls1738 signals exclusively through Ast-LsNRC1, and the Clade 2 sensor Ls2655 signals through both helpers. (**B**) Domain architecture of Ls2655, Ls0504, and Ls0504 chimeras carrying NB-domain interface stretch swaps from Ls2655. Colored bars represent the NB-domain interface stretches (S1–S4) required for helper activation. Ls2655 was selected as the donor for interface swaps due to its dual helper compatibility. Individual stretches (S1, S2, S3, S4) or combinations (S1/S2, S3/S4) from Ls2655 (highlighted) were introduced into the corresponding positions of Ls0504. Domain boundaries are indicated above. (**C**) Sequence alignment of NB domain stretches in Ls0504 and Ls2655. (**D**) Photo of representative WT *N. benthamiana* leaves showing cell death after co-expression of LsNRC0 or Ast-LsNRC1 with Ls2655 (WT or DV), Ls0504 (WT or DV), or Ls0504^DV^ chimeras carrying interface stretch swaps from Ls2655 (S1, S2, or S1/S2, as indicated). The experiments were repeated four times with at least six technical replicates per repeat, with similar results in all cases. Quantitative analysis of the cell death assays can be found in **Fig. S15**.

Strikingly, Ls0504^D436V^ chimeras carrying stretch 1 (S1), stretch 2 (S2), or the combined S1/S2 interface from Ls2655 gained the ability to initiate cell death via Ast-LsNRC1 while retaining signalling through LsNRC0 (**Fig. 8D, Fig. S15**). Among these, the S2 swap conferred the strongest gain of Ast-LsNRC1-dependent cell death. In contrast, none of the Ls1741^D489V^ chimeras acquired the ability to signal through LsNRC0, and several exhibited loss-of-function phenotypes, suggesting that certain interface swaps may disrupt stability or activation competence (**Fig. S16**). Together, these results suggest that the interface predicted for NRC network sensor-helper pairs across asterids is functionally relevant in at least two divergent species. Moreover, it shows that variation within this interface can modulate sensor-helper compatibility in this network.

### Two NB-domain point mutations enable Ls0504 to activate Ast-LsNRC1

Given that the Ls0504^DV–S2^ chimera displayed a gain of Ast-LsNRC1-dependent signalling, we next asked whether this expanded helper compatibility could be recapitulated by mutating individual residues within stretch 2. To this end, we identified all residues within stretch 2 that are polymorphic between Ls0504 and the promiscuous Clade 2 NRC-S Ls2655 and introduced each polymorphism individually into Ls0504, substituting the Ls0504 residue with the corresponding amino acid present at the equivalent position in Ls2655 (**Fig. 8C, Fig. S17A**). As in the chimera experiments, all point mutations were introduced in cis with the autoactive D to V substitution.

We tested the seven resulting Ls0504^DV^ point mutants for helper activation in cell death assays in leaves of *nrc2/3/4* CRISPR mutant *N. benthamiana* lines complemented with either LsNRC0 or Ast-LsNRC1. Two of these variants, Ls0504^L156Y-DV^ and Ls0504^D162H-DV^, gained the ability to activate Ast-LsNRC1 while retaining robust signalling through LsNRC0 (**Fig. S17A-B, Fig. S18**). In contrast, all remaining mutants behaved similarly to wild-type Ls0504^DV^ and remained restricted to signalling via LsNRC0 (**Fig. S16B, Fig. S17**). We generated an additional double mutant Ls0504^L156Y, D162H-DV^ carrying both mutations in cis. This variant also gained the capacity to signal through Ast-LsNRC1 and triggered a stronger cell death response than either single mutant alone (**Fig. S17, Fig. S18**).

Inspection of the AlphaFold 3-predicted complexes revealed that the two residues whose substitution expanded helper compatibility are positioned within the NB-domain interface and are predicted to form contacts with the LRR domain of both lettuce NRC helpers (**Fig. S17B**). Together, these results demonstrate that minimal sequence changes within the NB-domain interaction surface are sufficient to reprogram helper specificity in NRC-S, providing independent experimental support for the structural logic of NRC sensor–helper engagement predicted by AlphaFold 3.

## Discussion

In this study, we define a sensor–helper interface that is required for NRC helper activation in the NRC immune receptor network. Using AlphaFold 3–based protein structure modeling combined with extensive experimental validation, we predicted and validated a complex between the NRC-S NLR Rx and its helper NbNRC2a where the Rx NB domain makes extensive contacts with the NB and LRR domains of NbNRC2a. Comparative analyses of resting state and NbNRC2a-bound Rx suggests that NRC-S and NRC helper activation involve analogous structural transitions. Importantly, based on the predicted complex, binding of Rx is predicted to displace the NB domain of NbNRC2a, which could promote conformational rearrangements within the helper NB-ARC that are required for helper activation. Based on these findings, we propose a structurally informed working model for sensor-helper activation in the NRC network (**Fig. 9**).

**Figure 9.**
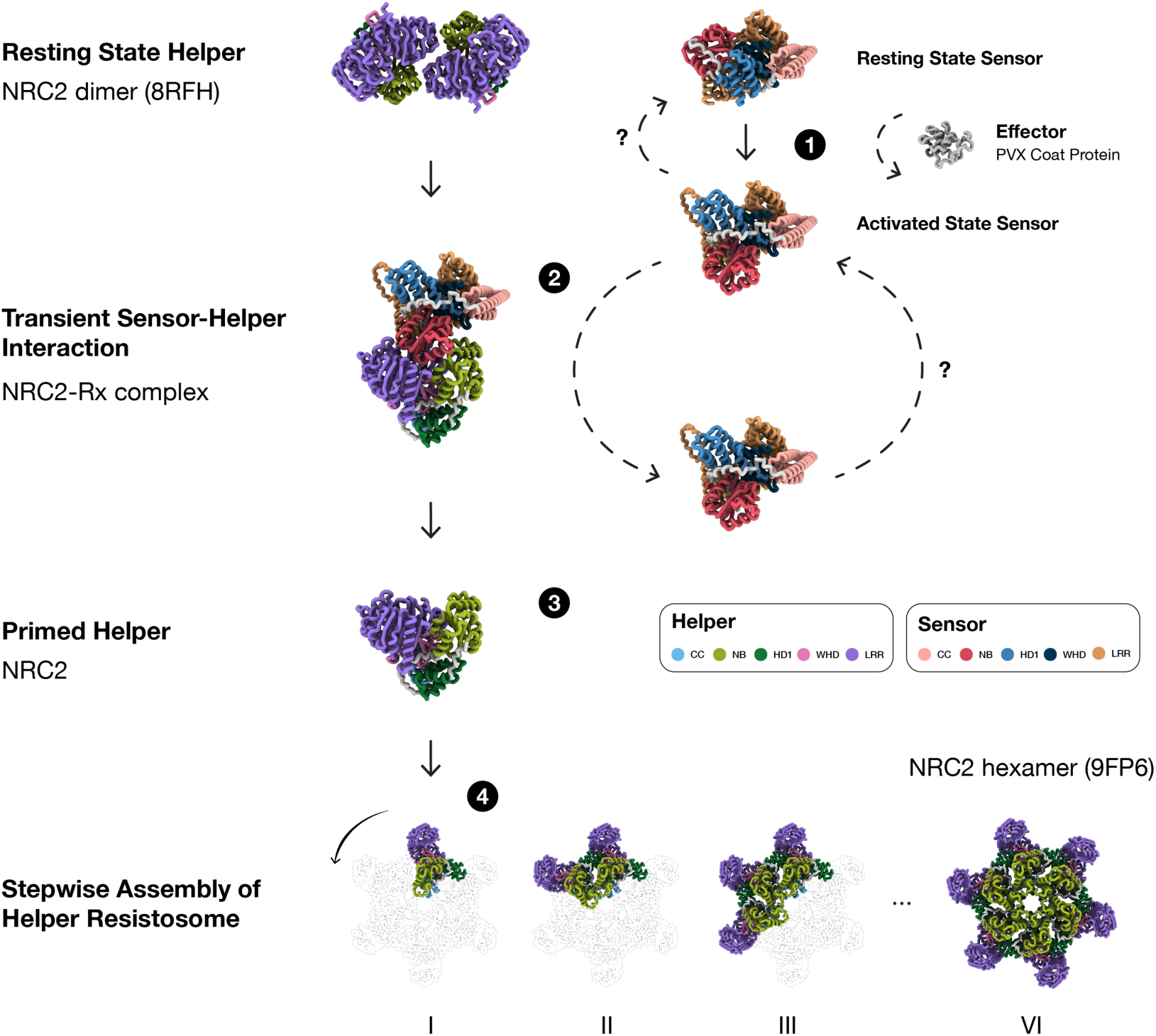
Structural working model for sensor-mediated NRC helper activation. Proposed stepwise activation-and-release mechanism for sensor–helper signaling in the NRC network, illustrated for the Rx–NbNRC2a pair. (1) In the resting state, NbNRC2a exists as an autoinhibited homodimer (PDB: 8RFH) and Rx adopts an inactive conformation in which the NB-domain helper interaction surface is occluded by intramolecular interactions. Upon recognition of the PVX coat protein, Rx undergoes activation-associated conformational rearrangements that expose the NB-domain interface. (2) The activated sensor transiently engages one protomer of the resting-state NbNRC2a homodimer through NB–NB and NB–LRR interactions. The predicted sensor binding site is sterically incompatible with the second homodimer protomer, suggesting that sensor engagement may promote helper homodimer dissociation. (3) Sensor binding displaces the NbNRC2a NB domain, promoting a primed helper conformation in which the NB domain is partially rotated outward relative to its position in the resting state. This primed intermediate would facilitate subsequent conformational rearrangements within the helper NB-ARC, including nucleotide exchange and NB–HD1 rotation, ultimately leading to sensor release. (4) The fully activated NbNRC2a protomers oligomerize stepwise into a hexameric resistosome (PDB: 9FP6), which accumulates at the plasma membrane to trigger the hypersensitive response and disease resistance. Protein domains are colored as indicated in the key. Question marks indicate steps that remain to be structurally characterized.

Extending beyond the Rx–NbNRC2a pair, AlphaFold 3 predictions across multiple asterid lineages revealed a conserved structural logic of sensor–helper interaction in NRC network pairs, defined by a shared binding mode and interaction topology despite extensive sequence divergence. Functional validation in the phylogenetically distant *Lactuca sativa* NRC network suggests that this predicted sensor-helper interface is functionally relevant in at least two asterid lineages. Importantly, structure-guided mutagenesis of NRC-S at this interface enabled expansion of downstream NRC helper compatibility, which suggests that this interface also encodes sensor-helper specificity. However, the extent to which the model presented in **Figure 9** is the ancestral mechanism of activation in the NRC network, and whether it applies to the extensive pairings of NRC-S and NRC helpers remains to be determined.

Beyond the functional characterization presented here, the interfaces identified by AlphaFold 3 and reported in this study align closely with prior mechanistic studies of sensor– helper communication in the NRC network (*19, 34*). We previously showed that the NB domains of multiple NRC-S are sufficient to activate NRC helpers and that these NB-domain truncations recapitulate the helper specificity profiles of their full-length counterparts (*19*). Consistent with these findings, AlphaFold 3 predicts that the NRC-S NB domain is the primary determinant of interaction with NRC helpers. Moreover, structure-guided swapping of helper interaction surfaces between lettuce NRC-S was sufficient to expand helper compatibility, indicating that helper specificity is largely encoded by NB-domain residues that directly contact helper NLRs. On the helper side, prior work demonstrated that mutations in the NB and LRR domains of NbNRC3 can enable compatibility with the oomycete resistance protein and NRC-S Rpi-blb2, implicating these domains in sensor-helper communication (*34*). Our model provides a unifying interpretation of these observations: NRC helpers engage activated NRC-S through direct interactions involving the NB and LRR domains of the helper (**Fig. 9**). Although these conclusions are partly based on structure prediction, their convergence with extensive functional evidence from this and previous studies strongly supports the biological relevance of the interface identified.

Based on the experimental support for our structure predictions, we expand the previously proposed activation-and-release model by incorporating a structural working model for NRC-S activation and communication with downstream NRC helpers (**Fig. 9**). In this model, autoinhibited resting-state NRC-S are unable to signal to helpers because the NB-domain interaction surface is occluded. Following pathogen effector recognition, rotation of the sensor NB–HD1 module exposes the helper interaction interface within the NB domain. The exposed sensor NB domain is then predicted to be recognized by NRC helpers through direct interactions with the helper NB and LRR domains, a process that may promote displacement of the helper NB domain. Notably, the predicted NRC-S binding geometry is sterically incompatible with the resting-state NbNRC2a homodimer, suggesting that sensor-helper interaction may trigger NRC helper homodimer dissociation (**Fig. S11**). Helper monomerization and NB-domain displacement were previously proposed to promote conformational rearrangements within the helper NB-ARC, potentially enabling nucleotide exchange, exposure of the helper oligomerization interface, and ultimately resistosome formation (*9, 11, 18, 19*).

Activated NRC-S like Rx trigger structural changes in NRC helpers analogous to those previously reported upon effector recognition in NLRs like ZAR1. Here, however, the activating ligand is not a pathogen-derived molecule but an endogenous danger signal: the exposed NB domain of the activated NRC-S. NRC helpers would thus represent an example of NLRs activated not by non-self (a pathogen-derived effector) but by modified self (a pathogen-activated upstream NRC-S). As proposed previously, NRCs could even be conceptualized as “guarding” the conformational status of their upstream NRC-S (*19*). This is analogous to the activation of the helper NLRs ADR1 and NRG1, where small immunogenic molecule-induced conformational changes in EDS1 heterodimers are perceived by the helper LRR, triggering conformational changes in the helpers that lead to activation and oligomerization (*35–39*). Together, these examples highlight the versatility of the LRR as a platform for immune activation, implicated in the perception of both pathogen-derived ligands and modified self.

Activated NRC resistosomes do not include sensor NLRs (*9, 18*), indicating that NRC-S such as Rx dissociate from helpers prior to resistosome assembly. How is the NRC-S ejected from the activation complex? In our model, the NRC-S engages the NRC helper through contacts with both the helper NB and LRR domains (**Fig. 9**). One possibility is that, once the helper NB-ARC adopts the active conformation, the approximately 180° rotation of the helper NB domain disrupts the dual interaction surface required for stable NRC-S binding, thereby destabilizing the sensor–helper complex. In addition, interactions between activated NRC helper protomers during oligomer assembly may occur with higher affinity than sensor–helper interactions, leading to ejection of the NRC-S from the complex. The precise interplay between NRC network sensor-helper and helper–helper binding affinities, interaction lifetimes, and activation strengths remains unknown; however, our findings raise the possibility that differences in these parameters may tune the efficiency and robustness of NRC helper activation by NRC-S across distinct sensor–helper pairs.

What are the evolutionary drivers of sensor/helper specificity in the NRC network? Functional experiments in lettuce demonstrate that NRC helper specificity can be determined by NRC-S NB amino acid stretches in the interaction interfaces. This suggests that specificity evolved through mutations in residues within a conserved structural framework rather than through fundamentally distinct interaction mechanisms. Narrow spectra of NRC network sensor-helper NLR specificity may arise as a consequence of co-evolution with pathogens, either to maintain effector recognition or to evade suppression by NLR-targeting effectors (*27, 30, 40*). Alternatively, selective advantages may be associated with limiting signaling routes, for example by reducing the risk of inappropriate activation or autoimmunity caused by incompatible sensor– helper interactions. Disentangling these paths will require the integration of evolutionary, functional, and structural analyses of NRC network diversification.

The ability to rewire compatibility between NRC-S and NRCs through structure-guided mutations highlights this interface as a promising target for engineering disease resistance. NRC- S with narrow NRC helper compatibility may be particularly vulnerable to pathogen suppression, as inhibition of a single helper could effectively disable immune signaling. Expanding helper NLR compatibility could therefore enhance the robustness of networked immune responses by allowing NRC-S to exploit the full signaling capacity of this network. At the same time, whether fully promiscuous sensor–helper communication would be beneficial or deleterious remains an open question, as unrestricted signaling could impose fitness costs or disrupt regulatory balance. Rational engineering of sensor–helper interfaces may enable tailored rewiring strategies that enhance immunity while preserving appropriate immune homeostasis. Notably, we achieved structure-guided engineering without experimentally solved structures, underscoring the potential of AlphaFold 3-based prediction to accelerate bioengineering of NLR immune receptors.

Our study highlights how structural modeling can advance our mechanistic understanding of NLR biology even in the absence of cryo-EM structures. AlphaFold 3 structural modeling allowed us to shed light on transient immune receptor interactions that proved recalcitrant to purification and experimental structure determination. Combining structure predictions with systematic loss- and gain-of-function validation allowed us to produce mechanistic and structural insights into NRC helper activation by NRC-S. Whether the sensor– helper interaction logic described here applies to other paired and networked NLRs beyond the expansive NRC network will be a fascinating topic to explore, possibly using AlphaFold modeling. It is becoming clear that multiple mechanisms of sensor–helper communication have evolved across plant NLR receptors (*35, 36, 41–43*). For instance, several NLR sensor-helper pairs form stable pre-activation complexes unlike NRC pairs (*41, 42, 44, 45*). To date, cryo-EM structures of heteromeric NLR complexes have remained elusive unlike the growing number of homomeric NLR structures obtained in presence or absence of pathogen effectors (*4*). Gaining structural insights into resting and activated sensor-helper NLR complexes will be essential for understanding the full diversity of NLR activation and signaling mechanisms.

## Data and materials availability

All data and code needed to evaluate and reproduce the results in the paper are present in the paper and/or the Supplementary Materials. All scripts used in this study are deposited at https://github.com/amiralito/Sensor_Helper_structural_logic. All AlphaFold 3 models have been deposited at https://doi.org/10.5281/zenodo.20085380 (*46*). The *N. benthamiana* lines, bacterial strains, and plasmids used in this study can be provided by S.K pending scientific review and a completed material transfer agreement through The Sainsbury Laboratory, Norwich, UK. Requests for *N. benthamiana* lines, bacterial strains, and plasmids should be submitted to.

## Acknowledgments

We thank all TSL support staff and Horticultural Services for preparing and providing plants and media. M.G. thanks La Colonia (Santa Fé, Argentina) for being the best helper network, La Pinta FC (Santa Fé, Argentina) for team spirit, and Chano for the energy. M.P.C. would like to thank Ben Seager (The Sainsbury Lab, Norwich, UK) for useful scientific discussions. M.P.C. would also like to thank Catriel Guerreiro and Ulises Guerriero (Buenos Aires, Argentina), for inspiration. M.G. and M.P.C. would like to thank the Argentinian public university and research system for the free, high-quality education and research training it has provided to many generations of scientists. A.T., M.G. and M.P.C. thank S. Bridge (The Sainsbury Lab, Norwich, UK) for holding things together.

## Author contributions

Conceptualization: A.T., S.K., M.P.C.

Methodology: A.T., M.G., R.F., M.P.C.

Validation: A.T., M.G., M.P.C.

Formal analysis: A.T., M.G., M.P.C.

Investigation: A.T., M.G., M.P.C.

Data curation: A.T., M.G., M.P.C.

Visualization: A.T., M.G., M.P.C.

Funding acquisition: S.K.

Project administration: A.T., S.K., M.P.C.

Supervision: S.K., M.P.C.

Writing—original draft: A.T., M.G., M.P.C.

Writing—review and editing: A.T., M.G., S.K., M.P.C.

## Materials and Methods

### Plant growth conditions

Wild-type and *nrc2/3/4* CRISPR mutant *N. benthamiana* lines were grown in a controlled environment growth chamber with a temperature range of 22 to 25 °C, humidity of 45% to 65% and a 16/8-hour light/dark cycle.

### Plasmid constructions

The Golden Gate Modular Cloning (MoClo) kit (*47*) and the MoClo plant parts kit (*48*) were used for cloning, and all vectors are from this kit unless specified otherwise. All Rx, Rx^NB^, NbNRC2a variants were generated via site directed mutagenesis (Genewiz) from previously published constructs (*18, 19*). Briefly, the original Rx and NbNRC2a constructs used for mutagenesis were cloned into the pJK001c acceptor with a 2x35S promoter (pICSL51288) and a 35S terminator (pICSL41414) with a C-terminal 3xFLAG tag (pICSL5007). The original Rx^NB^ construct used for mutagenesis was cloned into the pJK268c acceptor with a 2x35S promoter (pICSL51288) and a 35S terminator (pICSL41414) with a C-terminal eGFP tag (pICSL50034). Note that the pJK268c acceptor features a P19 expression cassette in the same T-DNA as the gene of interest to boost expression. Ls0504, Ls1734, Ls2655, LsNRC0 and Ast-LsNRC1 constructs were synthesized (Genewiz) and cloned into the pJK001c acceptor with a 2x35S promoter (pICSL51288) and a 35S terminator (pICSL41414) using a C-terminal 4xMyc (pICSL5010) for all sensors or a 3xFLAG tag (pICSL5007) for the helpers. eGFP, SlNRC0- 3xFLAG and PVX CP-eGFP constructs were all published previously. Cloning design and sequence analysis were done using Geneious Prime (v2021.2.2; https://www.geneious.com).

### Transient protein expression in *N. benthamiana* by agroinfiltration for cell death assays

NLR immune and effectors of interest were transiently expressed according to previously described methods (*49*). Briefly, leaves from 4–5-week-old plants were infiltrated with suspensions of *A. tumefaciens* GV3101 pM90 strains transformed with expression vectors coding for different proteins indicated. Final OD_600_ of all *A. tumefaciens* suspensions were adjusted in infiltration buffer (10 mM MES, 10 mM MgCl_2,_ and 150 µM acetosyringone (pH 5.6)). Final OD_600_ used was 0.3 for all Rx, RxNB or NbNRC2a variants used and 0.2 for PVX CP-eGFP or eGFP. Leaves were imaged at 5 or 7 days using a camera for brightield or an ImageQuant 800 imaging system for fluorescent images using the Cy3 channel (excitation: 535 nm, emission: Cy3 UV filter, exposure time: 3 seconds). Cell death was scored using a modified 0–7 scale at 3–5 days post agroinfiltration as described previously (*50*).

### Extraction of total proteins for BN-PAGE and SDS-PAGE assays

Four to five-week-old plants were agroinfiltrated as described using the OD_600_ described above with constructs of interest and leaf tissue was collected 3 days post agroinfiltration. Leaf tissue was ground using a GenoGrinder tissue homogenizer and total protein was subsequently extracted and homogenized extraction buffer. Total proteins were extracted in the GTEN buffer [10% v/v glycerol, 50 mM tris-HCl (pH 7.5), 1 mM EDTA, and 50 mM NaCl], supplemented with 10 mM DTT, 1× protease inhibitor cocktail (Sigma-Aldrich), and 0.2% v/v IGEPAL (Sigma-Aldrich), and incubated on ice for 10 min. Samples were incubated in extraction buffer on ice for 15 minutes with short vortex mixing at every 2 minutes. Following incubation, samples were centrifuged at 5,000 x*g* for 15 minutes and the supernatant was used for SDS- PAGE assays.

### SDS-PAGE assays

For SDS-PAGE, samples were diluted in SDS loading dye and denatured at 72 °C for 10 minutes. Denatured samples were spun down at 5,000 x*g* for 3 minutes and supernatant was run on 4%-20% Bio-Rad 4%-20% Mini-PROTEAN TGX gradient gels alongside a PageRuler Plus prestained protein ladder (Thermo Scientific). The proteins were then transferred to polyvinylidene difluoride membranes using Trans-Blot Turbo Transfer Buffer using a Trans-Blot Turbo Transfer System (Bio-Rad) as per the manufacturer’s instructions. Membranes were immunoblotted as described below.

### Immunoblotting and detection of BN-PAGE and SDS-PAGE assays

Blotted membranes were blocked with 5% milk in Tris-buffered saline plus 0.01% Tween 20 (TBS-T) for an hour at room temperature and subsequently incubated with desired antibodies at 4 °C overnight. Antibodies used were anti-GFP (B-2) HRP (Santa Cruz Biotechnology) and anti-FLAG (M2) HRP (Sigma), anti-Myc (Y69) HRP (Sigma), and anti-HA (3F10) HRP (Sigma), all used in a 1:5000 dilution in 5% milk in TBS-T. To visualize proteins, we used Pierce ECL Western (32106, Thermo Fisher Scientific), supplementing with up to 10% SuperSignal West Femto Maximum Sensitivity Substrate (34095, Thermo Fisher Scientific) when necessary. Membrane imaging was carried out with an ImageQuant 800 luminescent imager (GE Healthcare Life Sciences, Piscataway, NJ). Rubisco loading control was stained using Ponceau S (Sigma) or Ponceau 4R (AG Barr).

### Phylogenetic analysis

We first extracted 91,366 NLR proteins from a dataset of 230 superasterid genomes annotated by Helixer (*51, 52*). Based on NLRtracker output, 79,206 NLRs with “CNL”, “CNLO”, “CN”, “OCNL”, “CONL”, “NL”, “NLO”, “ONL”, “BCNL”, “BNL”, “BNLO”, “BBNL”, “BCN”, “BCCNL”, “BBCNL”, “BOCNL”, “RNL”, “TN”, “TNL”, “TNLO”, and “TNLJ” domain architectures were retained (*10*). Redundant sequences were removed using CD-HIT v4.8.1, resulting in 77,828 non-redundant NLR proteins (*53*). Sequences with NB-ARC domains shorter than 250 or longer than 400 amino acids were removed to avoid truncated or misannotated proteins, yielding a final set of 67,438 NLRs. During manual inspection, we found that the previously described Beta vulgaris NRC-S was missing from the Helixer annotation (*7*). This sensor NLR was therefore manually added from the RefSeq annotation set (XP_048500768.1) (*54, 55*). NB-ARC domains from this dataset were aligned with the RefPlantNLR reference set using FAMSA v2.4.1 (*56*), and the resulting alignment was used to construct a phylogenetic tree with FastTree v2.1.11 using the -lg option (*57*). Based on the presence of reference NRC helper and NRC-S sequences, we defined and extracted the NRC superclade, which contained 24,676 sequences. We then reconstructed a phylogenetic tree from the extracted NRC superclade using the same approach. The final tree was rooted on the NRC helper clade and visualized using iTOL v6 (*58*).

### Alphafold modeling and structural analysis

All models were generated using AlphaFold 3 web server (https://alphafoldserver.com/) (*20–22*). Solanaceae sensor–helper complexes were modeled 10 times using seeds 1–10, whereas complexes from the remaining lineages were modeled using seeds 1–5. Monomeric structures were predicted with ADP to simulate the NLR resting state conformation. Structural alignments were generated using ChimeraX-1.10 matchmaker command (*59*). Contacts were extracted from the resulting models and visualized using custom Python scripts. All structures were visualized with ChimeraX. All scripts are deposited at https://github.com/amiralito/Sensor_Helper_structural_logic. All AlphaFold 3 models have been deposited at https://doi.org/10.5281/zenodo.20085380 (*46*).

