## Supplementary Figures and Legends for "A conserved structural logic underlies sensor–helper NLR communication in the NRC immune receptor network"

### Supplementary Data

**Data S1.** Sequence and metadata of the modeled NRC, NRC-S, and non-NRC NLRs.

### Supplementary Figures

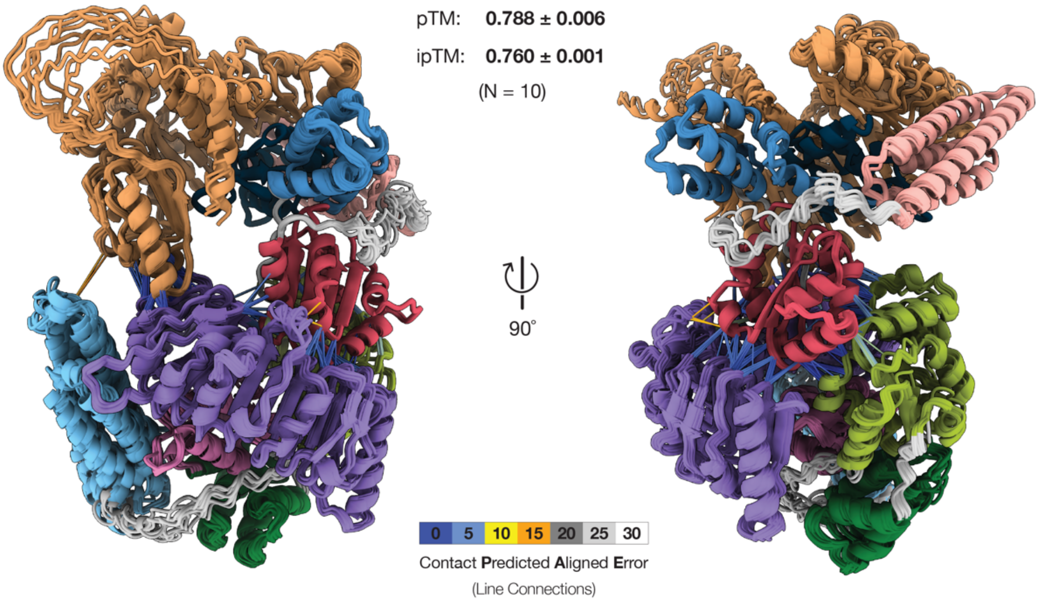


**Figure S1. AlphaFold 3 predicts a high confidence complex between the sensor Rx and its helper NbNRC2a across 10 replicates.**

Selected views of the predicted model of Rx in complex with NbNRC2a. The predictions were run in 10 replicates with seeds 1 to 10. An overlay of the 10 models is shown. Structures of Rx and NbNRC2a are colored based on the schematic representation of the domain architectures of each protein. Average and SD values for ipTM and pTM (N = 10) are shown on the right.

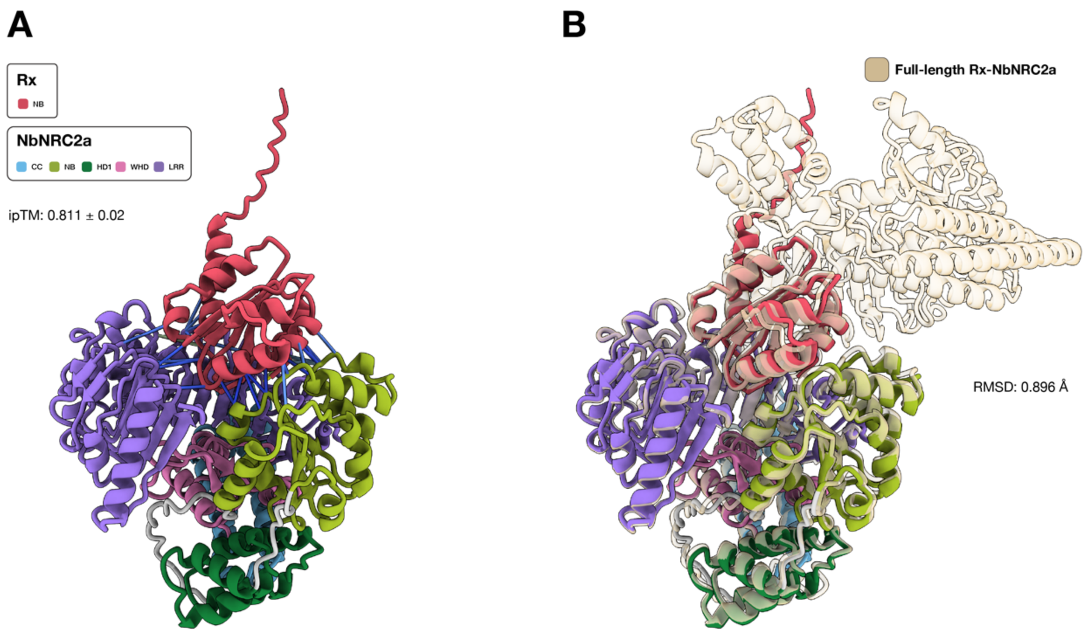


**Figure S2. AlphaFold 3 predicts a high confidence complex between the sensor Rx$^{NB}$ and its helper NbNRC2a across 10 replicates, aligning well with the full-length Rx-NbNRC2a complex.**

Selected view of the predicted model of **(A)** Rx$^{NB}$ and **(B)** full-length Rx in complex with NbNRC2a. The predictions were run in 10 replicates with seeds 1 to 10. Structures of Rx and NbNRC2a are colored based on the schematic representation of the domain architectures of each protein. Average and SD values for ipTM and pTM (N = 10) are shown on the top left. The two complexes aligned with a Root Mean Square Deviation (RMSD) of 0.896 Å.

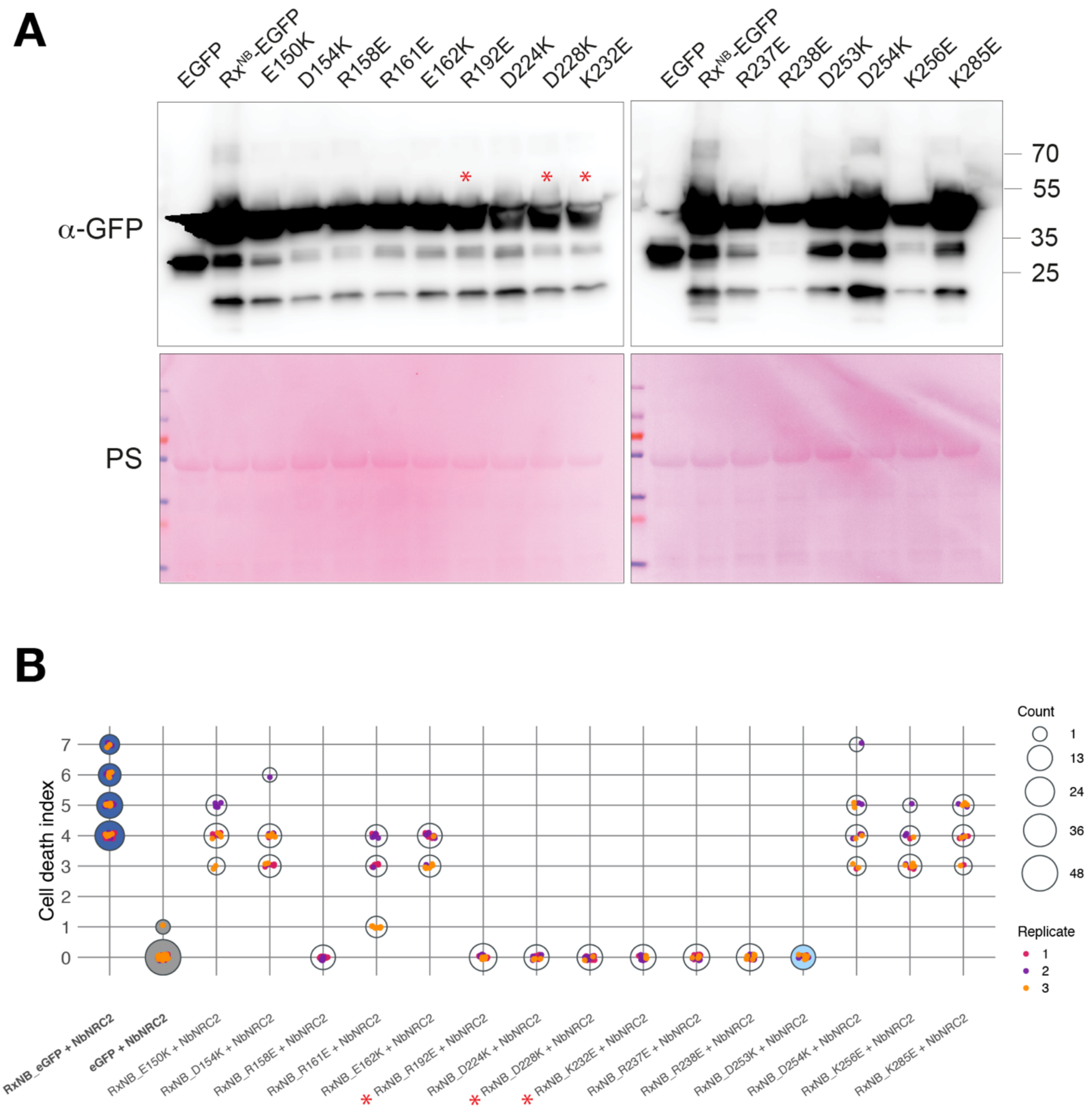


**Figure S3: Rx$^{NB}$ loss-of-function variants accumulate *in planta*.**

(**A**) Protein extracts from leaves of *nrc2/3/4* KO *N. benthamiana* expressing proteins of interest were run on SDS-PAGE assays and immunoblotted with the appropriate antisera labelled on the left. Approximate molecular weights (kDa) of the proteins are shown on the right. Rubisco loading control was carried out using Ponceau staining (PS). Red asterisks indicate the four variants which lost the capacity to signal through all NRC helpers tested. Experiment was repeated two times with similar results. (**B**) Quantitative analysis of cell death assays shown in **Fig. 3C**. Cell death was scored using a modified 0–7 scale at 3–5 days post agroinfiltration (*50*). Dot size is proportional to the number of samples with the same score (Count). Data represent three biological replicates.

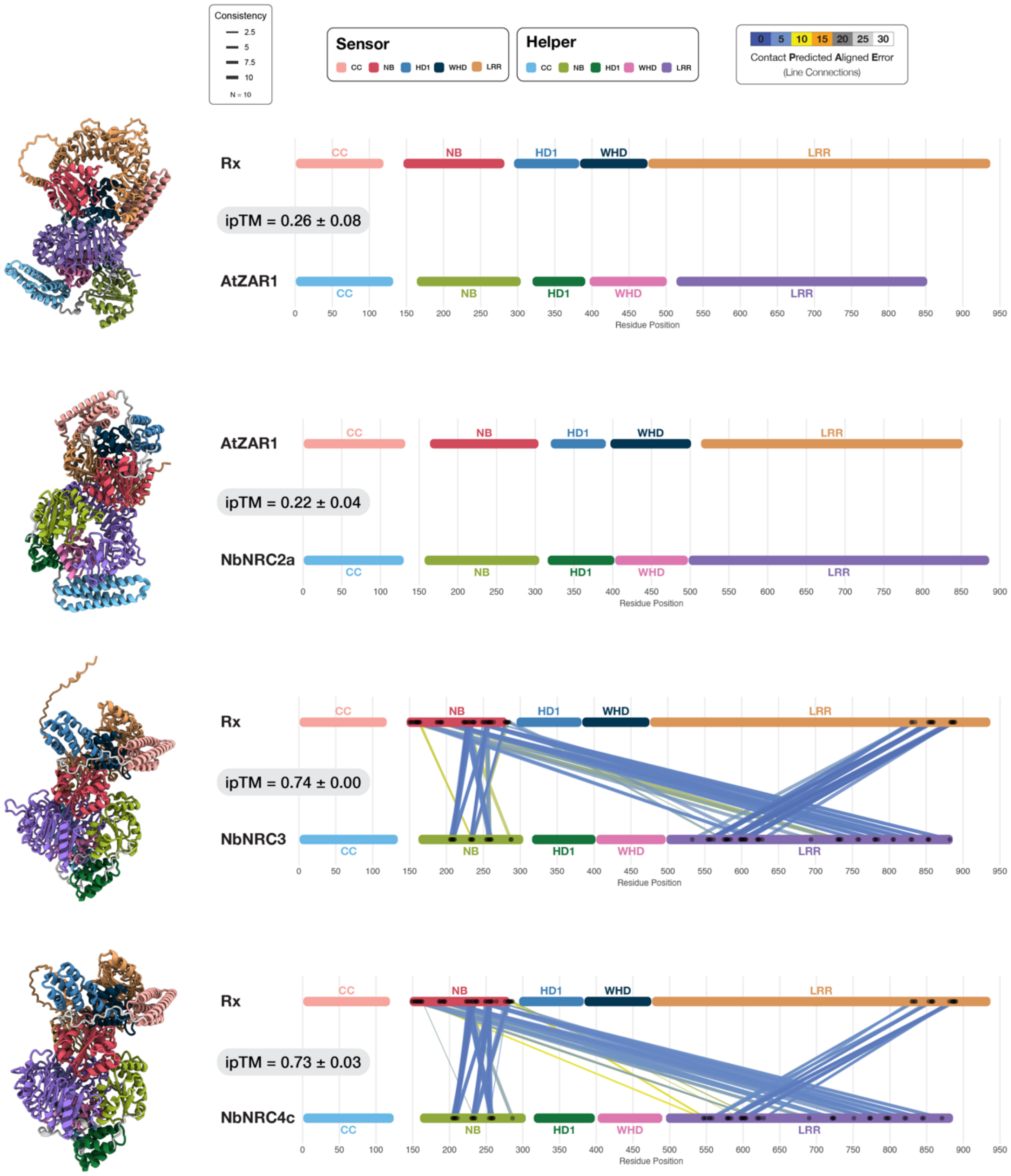


**Figure S4. The predicted Rx and NbNRC3 and NbNRC4c complexes involve NB-NB, NB-LRR, and LRR-LRR interactions.**

Schematic representation of the domain architectures of Rx, NbNRC2a, NbNRC3, NbNRC4c, and AtZAR1. Lines indicate contacts between the two proteins at the connected positions, as predicted by AlphaFold 3. AtZAR1 was used as helper control for predicting complexes with Rx, and as sensor control for predicting complexes with NbNRC2a. The color of the lines indicates the contact Predicted Aligned Error for that interaction and the line thickness indicates the consistency of that contact across all 10 modeling replicates.

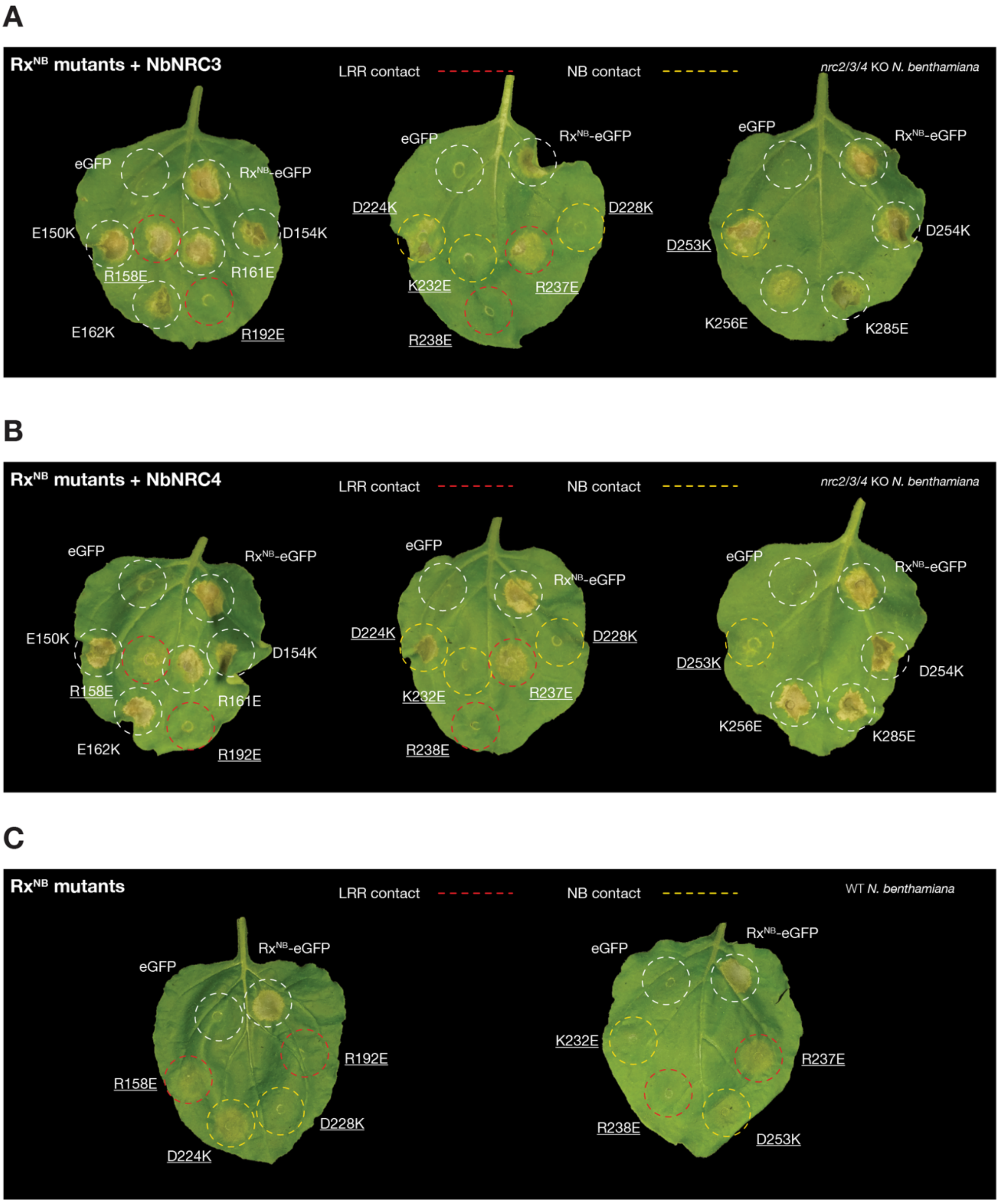


**Figure S5. Four Rx$^{NB}$ interface residues are required for activation of NbNRC2a, NbNRC3, and NbNRC4c.**

(**A**) Representative *nrc2/3/4* KO *N. benthamiana* leaf images showing cell death after co-expression of NbNRC3 with different Rx$^{NB}$-eGFP variants. Wild-type Rx$^{NB}$-eGFP and free eGFP were co-expressed with NbNRC3 as positive and negative controls for cell death, respectively. (**B**) Representative *nrc2/3/4* KO *N. benthamiana* leaf images showing cell death after co-expression of NbNRC4 with the indicated Rx$^{NB}$-eGFP variants. Wild-type Rx$^{NB}$-eGFP and free eGFP were co-expressed with NbNRC4 as positive and negative controls for cell death, respectively. For both panels, mutations that led to loss of helper activation are outlined with dashed circles; the color of the dotted outline indicates whether the corresponding Rx residue was predicted to make contacts with the helper NB (yellow) or LRR (red) domain. (**C**) Representative WT *N. benthamiana* leaf images showing cell death after expression of wild-type Rx[NB]-eGFP (positive control), eGFP (negative control), or the indicated Rx[NB]-eGFP point mutants in the absence of a co-expressed helper. All experiments were repeated three times with at least six technical replicates per repeat, with similar results in all cases.

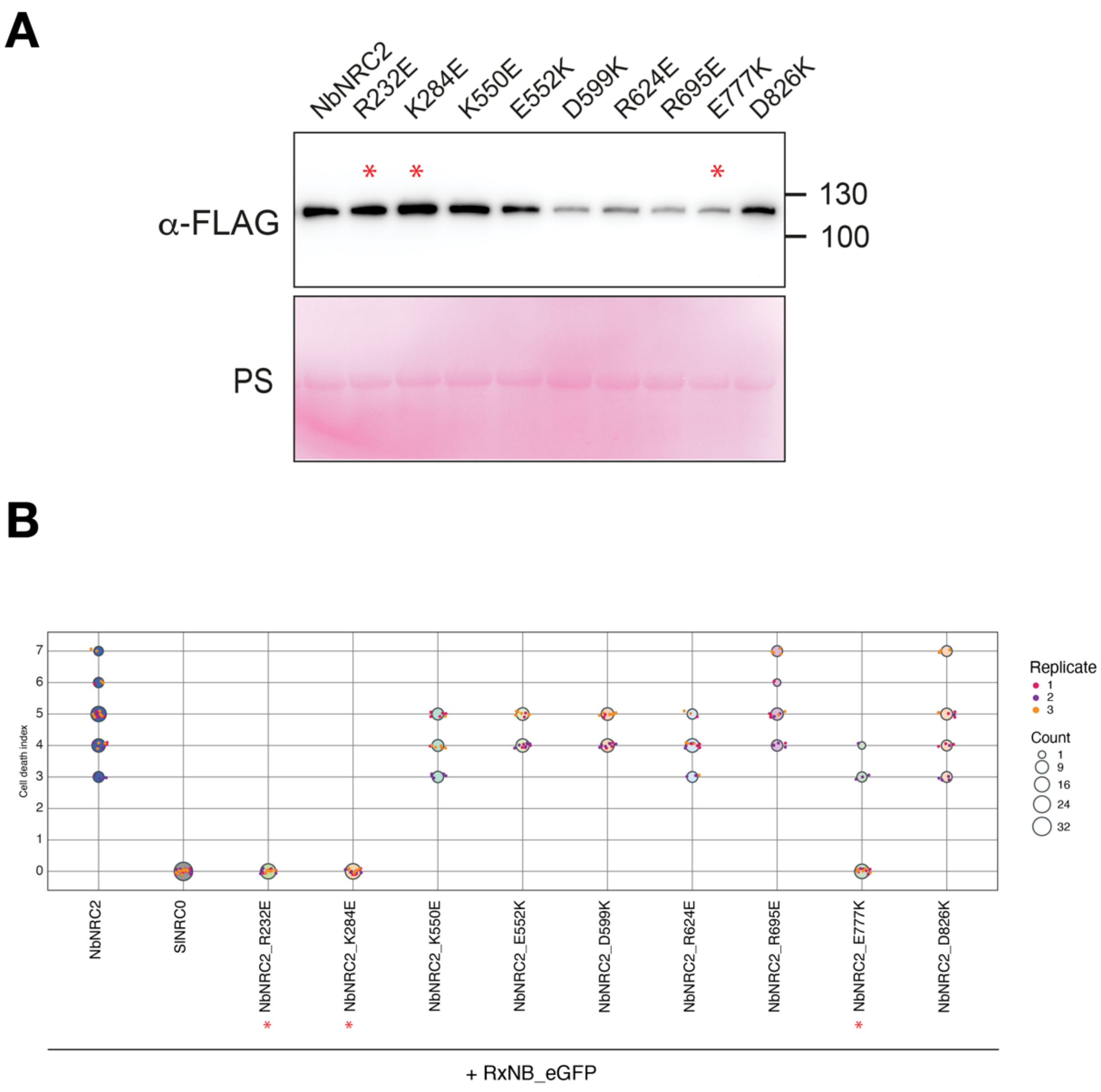


**Figure S6. NbNRC2a loss-of-function mutants accumulate *in planta*.**

(**A**) Protein extracts from leaves of *nrc2/3/4* KO *N. benthamiana* expressing proteins of interest were run on SDS-PAGE assays and immunoblotted with the appropriate antisera labelled on the left. Approximate molecular weights (kDa) of the proteins are shown on the right. Rubisco loading control was carried out using Ponceau staining (PS). Red asterisks indicate the three NbNRC2a variants that lost the capacity to be activated by Rx[NB]. Experiment was repeated two times with similar results. (**B**) Quantitative analysis of cell death assays shown in **Fig. 4B**. Cell death was scored using a modified 0–7 scale at 3–5 days post agroinfiltration (*50*). Dot size is proportional to the number of samples with the same score (Count). Data represent three biological replicates. Variants that lost the capacity to support $Rx^{NB}$-dependent cell death are indicated with red asterisks below the x-axis, consistent with panel (A).

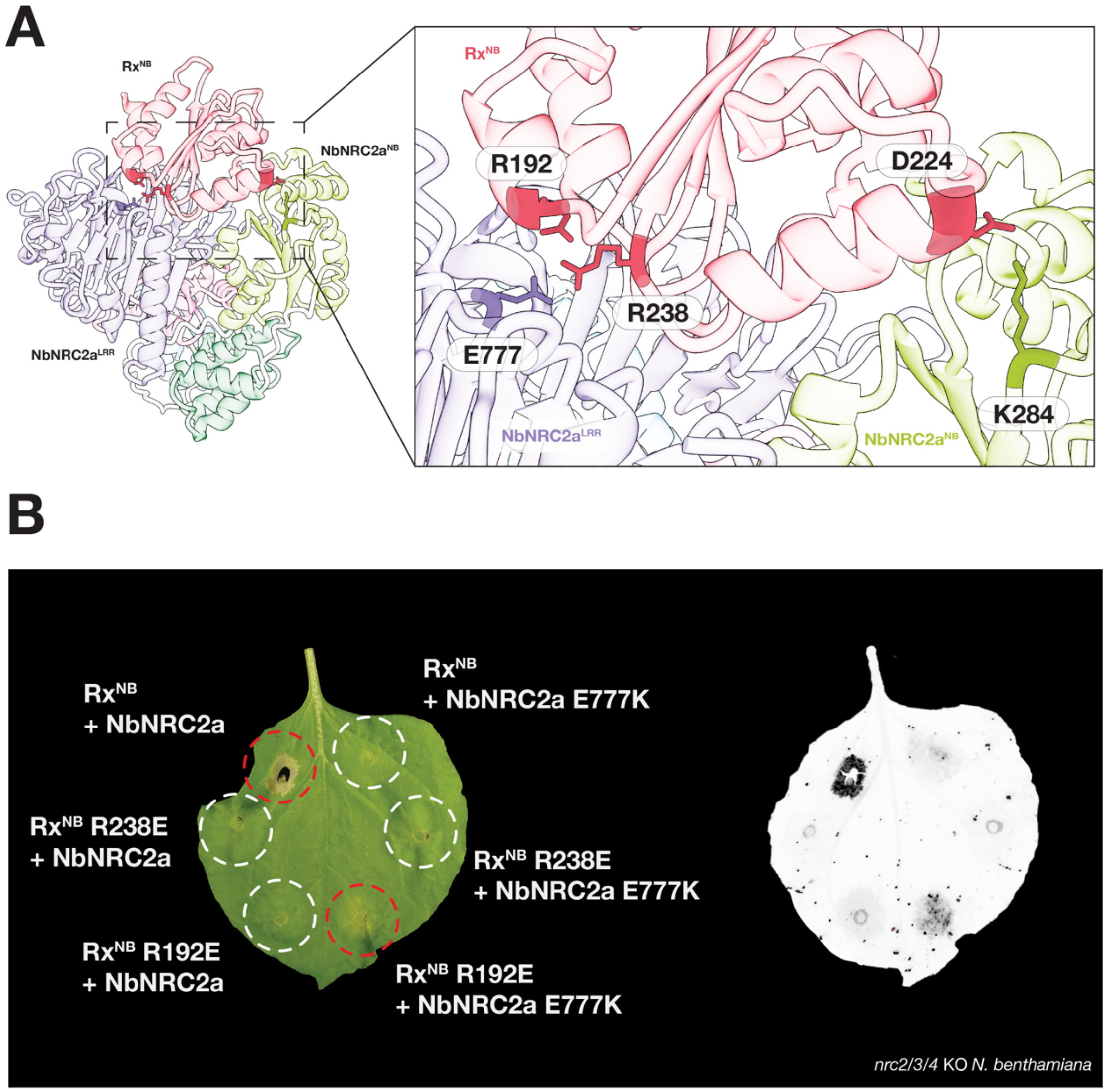


**Figure S7. Additional reciprocal charge swaps weakly restore Rx$^{NB}$–NbNRC2a signaling.**

(**A**) AlphaFold 3-predicted salt bridges across the Rx$^{NB}$–NbNRC2a interface highlighting the residue pairs targeted for reciprocal charge-swap mutagenesis. (**B**) Representative *nrc2/3/4* KO *N. benthamiana* leaf images (bright-field and UV) showing cell death following co-expression of Rx$^{NB}$ with NbNRC2a (positive control), and the additional Rx$^{NB}$ or NbNRC2a single charge-swap mutants paired with the corresponding WT partner, or matched Rx$^{NB}$/NbNRC2a reciprocal charge-swap combinations. The experiment was repeated three times with at least six technical replicates per repeat, with similar results.

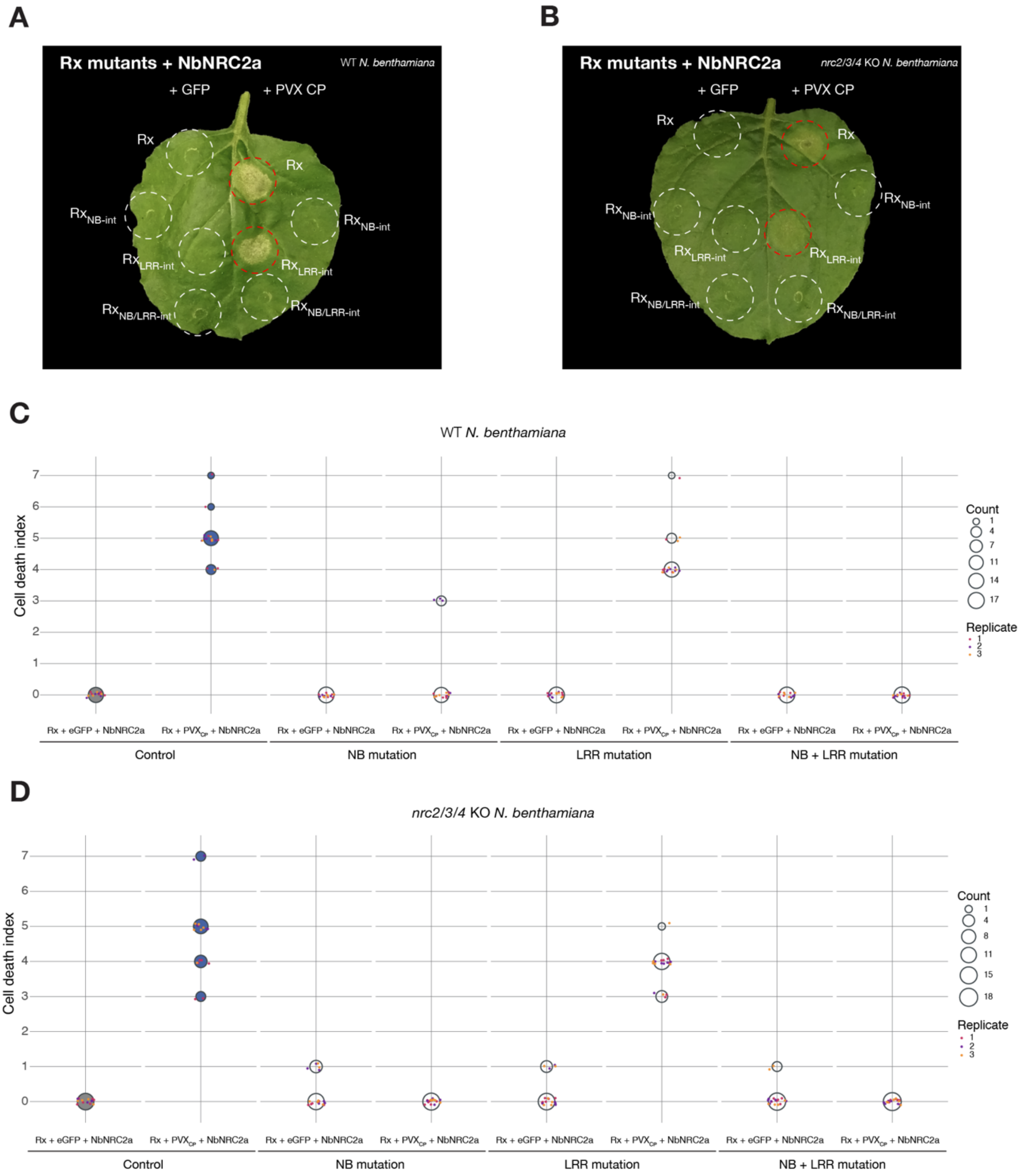


**Figure S8. NB-domain interactions, but not the LRR–LRR interface, are required for helper activation by full-length Rx.**

(**A**) Representative WT N. benthamiana and (**B**) nrc2/3/4 KO N. benthamiana leaf images showing cell death after co-expression of NbNRC2a with full-length Rx or the indicated Rx interface variants ($Rx_{NB\text{-}int}$, $Rx_{LRR\text{-}int}$, or $Rx_{NB/LRR\text{-}int}$) in the presence of eGFP or PVX CP. Wild-type Rx co-expressed with eGFP and PVX CP served as negative and positive controls for NbNRC2a-dependent cell death, respectively. Dashed circles indicate infiltration zones; red outlines denote zones displaying cell death. The experiments were repeated three times with at least six technical replicates per repeat, with similar results in all cases. (**C, D**) Quantitative analysis of cell death assays shown in (**A**) and (**B**), respectively. Cell death was scored using a modified 0–7 scale at 3–5 days post agroinfiltration (*50*). Dot size is proportional to the number of samples with the same score (Count). Data represent three biological replicates.

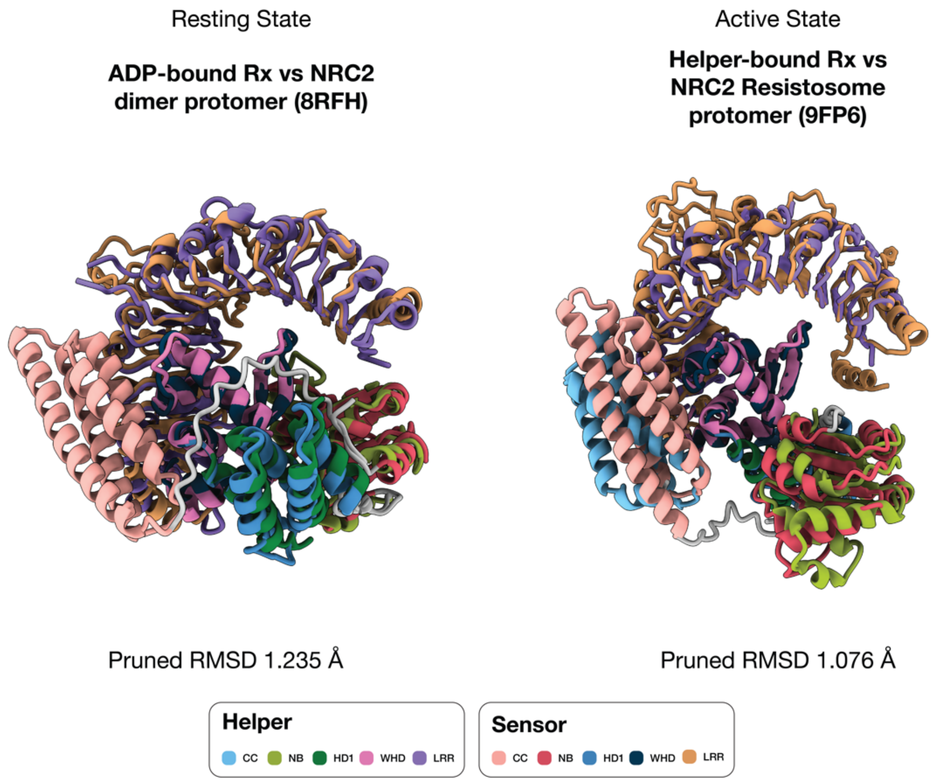


**Figure S9. Predicted resting and NbNRC2a-bound Rx conformations mirror structural transitions observed during helper activation.**

(Left) Structural superposition of AlphaFold 3-predicted resting-state Rx (ADP-bound) with a protomer from the experimentally determined resting-state NbNRC2a homodimer (PDB: 8RFH). Pruned RMSD = 1.235 Å. (Right) Structural superposition of AlphaFold 3-predicted helper-bound Rx (from the Rx–NbNRC2a complex prediction) with an activated NbNRC2a protomer from the experimentally determined hexameric resistosome (PDB: 9FP6). Pruned RMSD = 1.076 Å. Structures are colored by domain as indicated in the color key below.

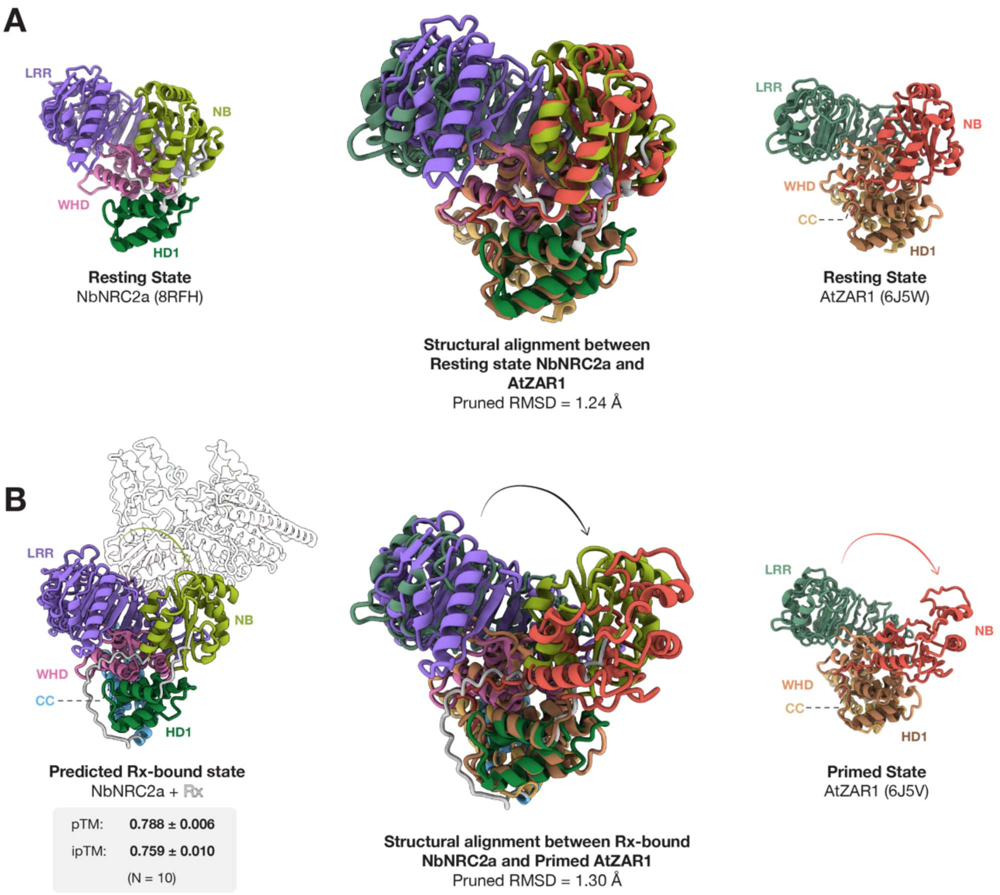


**Figure S10. The predicted Rx-bound NbNRC2a conformation resembles the primed state of AtZAR1.**

**(A)** Structural alignment of resting-state NbNRC2a (PDB: 8RFH, left) with resting-state AtZAR1 (PDB: 6J5V, right). Structures are colored by domain. The center panel shows the superposition of the two structures (pruned RMSD = 1.24 Å), illustrating the conserved overall domain arrangement between resting-state NbNRC2a and AtZAR1. **(B)** Structural alignment of the AlphaFold 3-predicted Rx-bound NbNRC2a conformation (left) with the primed state of AtZAR1 (PDB: 6J5V, right). Mean pTM and ipTM values (± SD, N = 10) for the Rx–NbNRC2a complex prediction are indicated. The center panel shows the superposition of the two structures (pruned RMSD = 1.30 Å), revealing that the NbNRC2a conformation adopted upon predicted Rx binding closely resembles the experimentally determined primed intermediate of AtZAR1, in which the NB domain is partially rotated outward relative to the resting state.

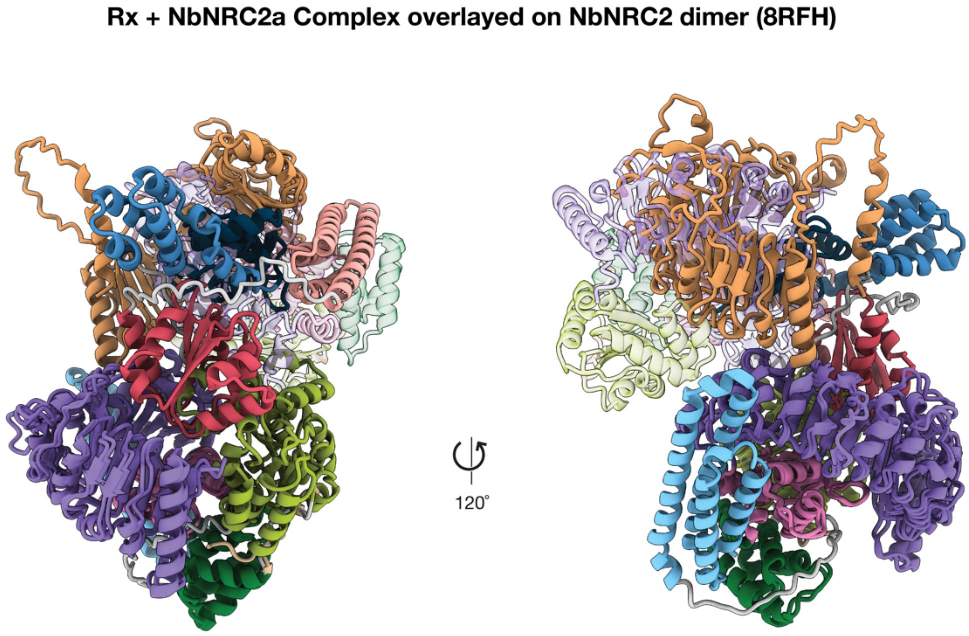


**Figure S11. The predicted Rx–NbNRC2a complex is sterically incompatible with the resting-state NbNRC2a homodimer.**

Two views (related by a 120° rotation) of the AlphaFold 3-predicted Rx–NbNRC2a complex superimposed onto the experimentally determined resting-state NbNRC2a homodimer (PDB: 8RFH). The predicted complex (Rx and NbNRC2a colored by domain as in **Fig. 1**) was aligned to one protomer of the NbNRC2a homodimer; the second homodimer protomer is shown in faded colors. The superposition reveals extensive steric clashes between the predicted position of Rx and the second NbNRC2a protomer.

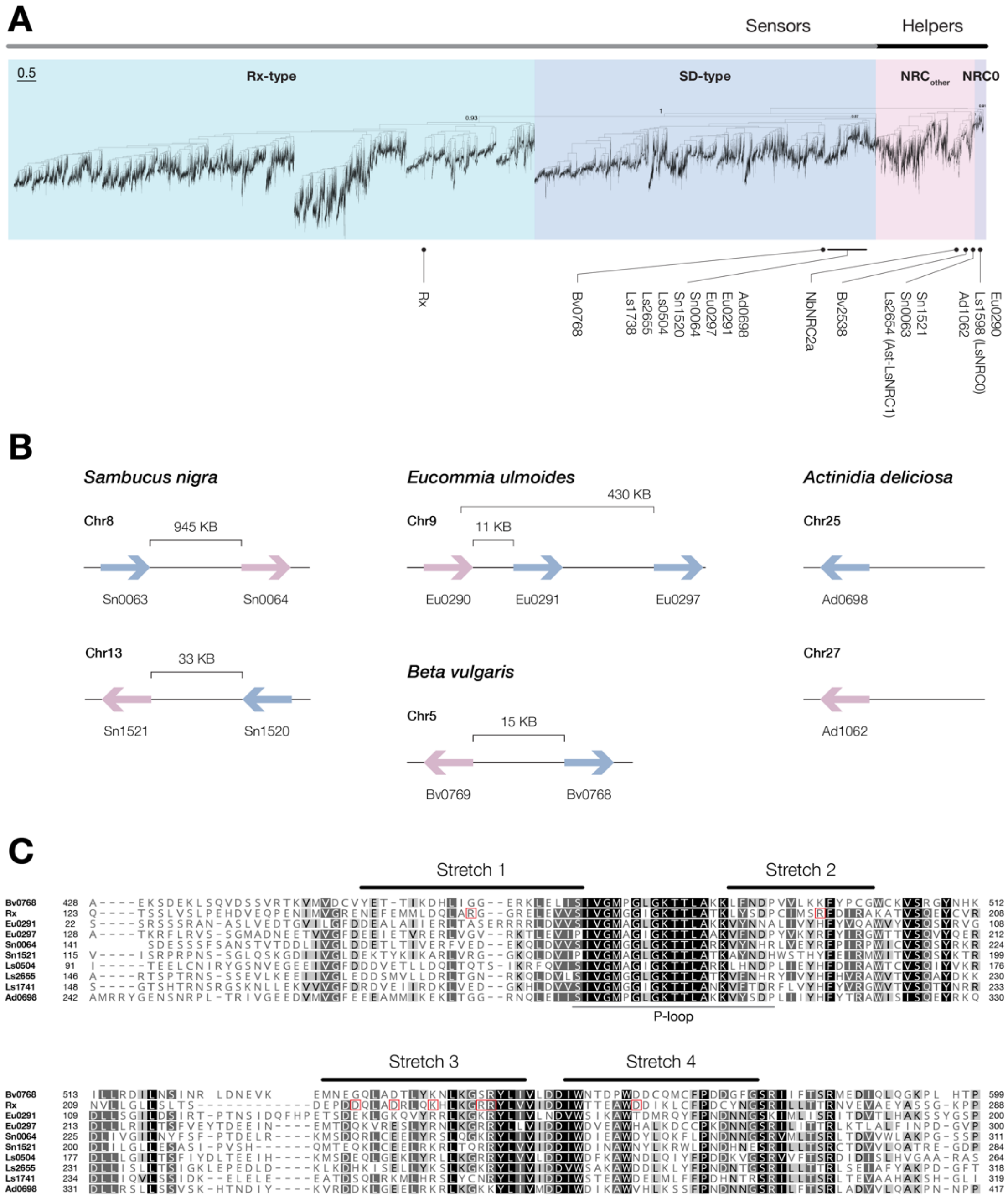


**Figure S12. Representative NRC sensor-helper pairs across asterids.**

**(A)** Phylogenetic tree of 24,668 NB-ARC sequences from NRC-S and helper sequences, together with 49 reference NRC helper and NRC-S sequences from RefPlantNLR (*10*). The tree is colored by sensor and helper classes. Representative NRC sensors and helpers used for AlphaFold 3 modeling are shown below the tree with their approximate phylogenetic clade. **(B)** Genomic location of the representative sensor-helper pairs from *Eucommia ulmoides*, *Sambucus nigra*, *Actinidia deliciosa*, *Lactuca sativa*, and *Beta vulgaris*. Schematics are not to scale. (C) Multiple sequence alignment of NB domains from the studied NRC-S. Sensor-Helper interface stretches are annotated based on $Rx^{NB}$. $Rx^{NB}$ mutants with loss-of-function phenotype are highlighted with red squares.

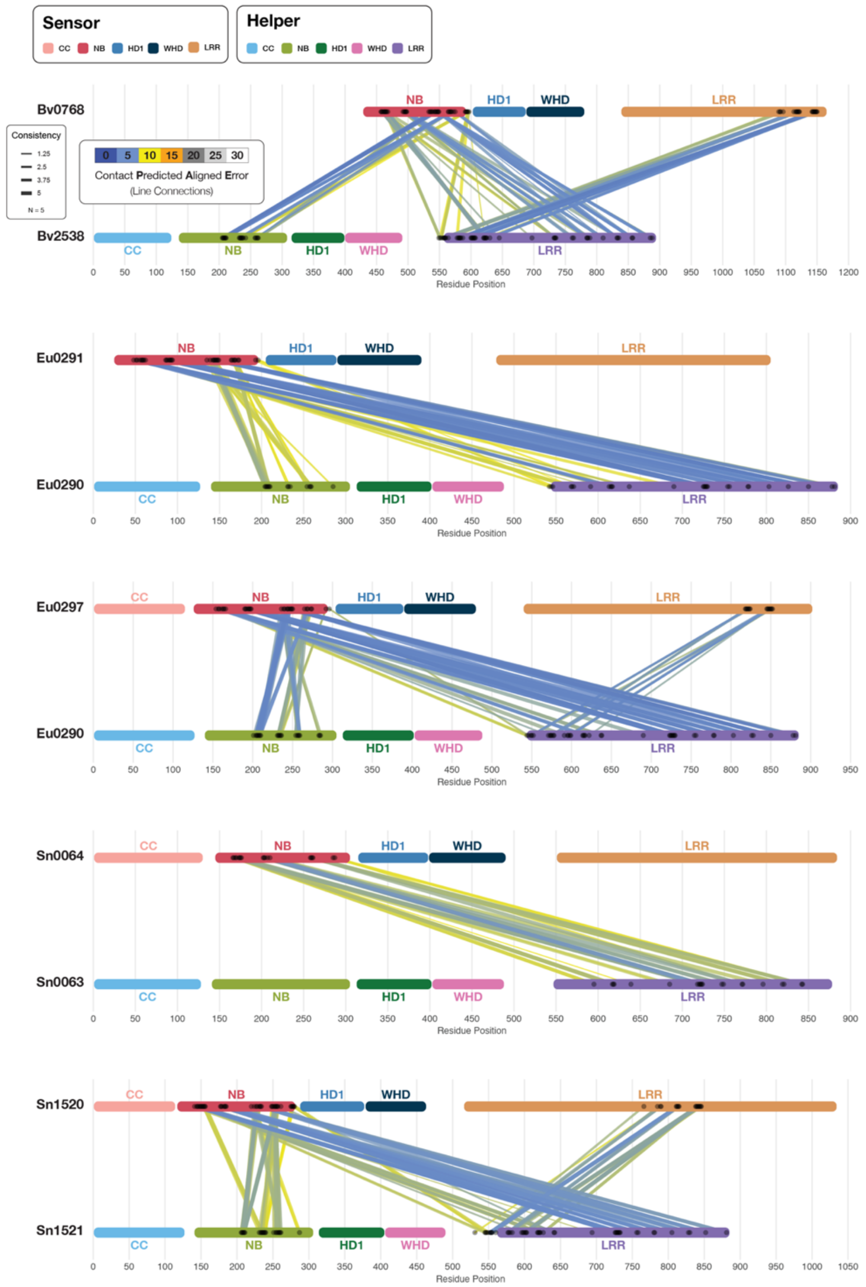


**Figure S13. Predicted sensor-helper interfaces in representative pairs from *Eucommia ulmoides*, *Sambucus nigra*, and *Beta vulgaris*.**

Schematic representation of the domain architectures of sensor-helper pairs from *Eucommia ulmoides*, *Sambucus nigra,* and *Beta vulgaris*. Lines indicate contacts between the two proteins at the connected positions, as predicted by AlphaFold 3. The color of the lines indicates the contact Predicted Aligned Error for that interaction and the line thickness indicates the consistency of that contact across all 5 modeling replicates.

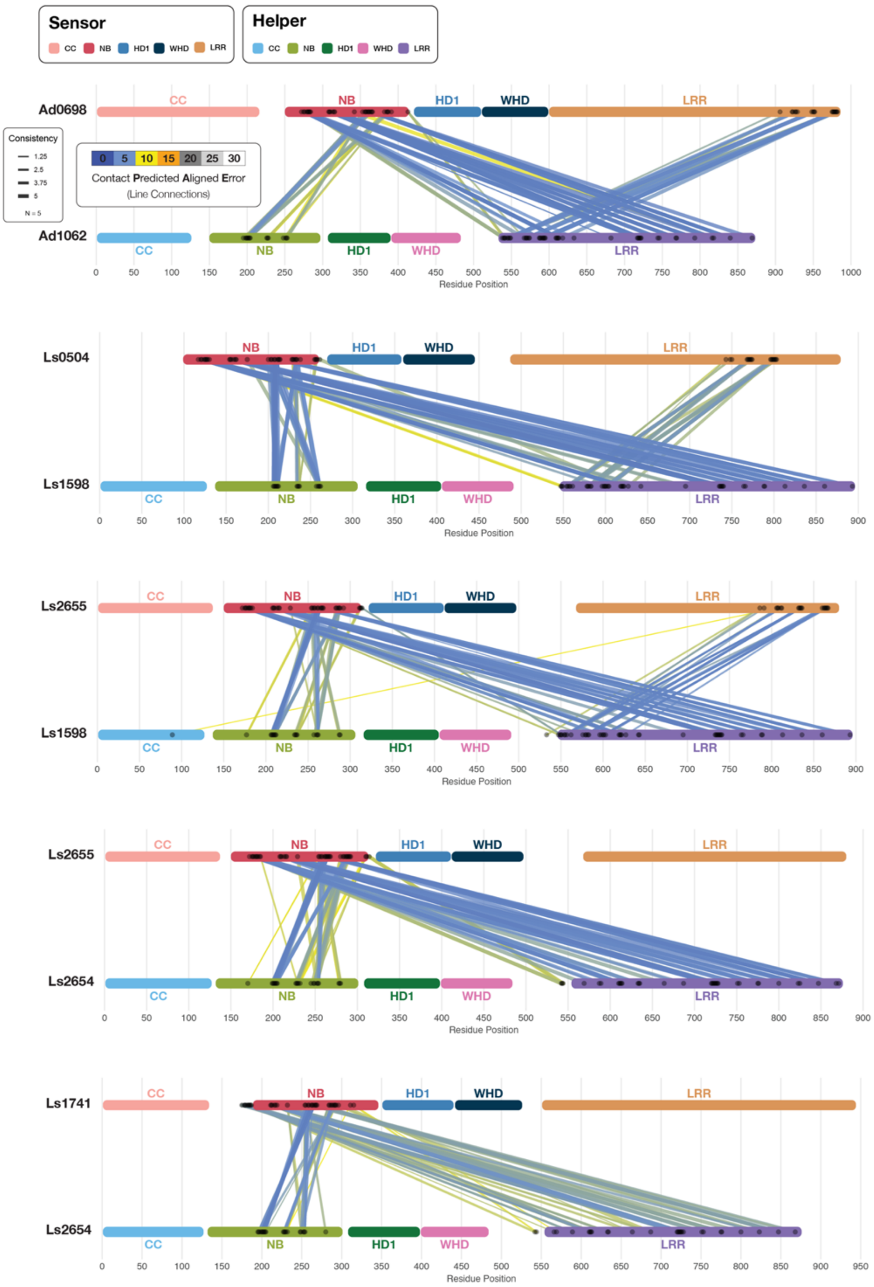


**Figure S14. Predicted sensor-helper interfaces in representative pairs from *Actinidia deliciosa* and *Lactuca sativa.***

Schematic representation of the domain architectures of sensor-helper pairs from *Actinidia deliciosa* and *Lactuca sativa*. Lines indicate contacts between the two proteins at the connected positions, as predicted by AlphaFold 3. The color of the lines indicates the contact Predicted Aligned Error for that interaction and the line thickness indicates the consistency of that contact across all 5 modeling replicates.

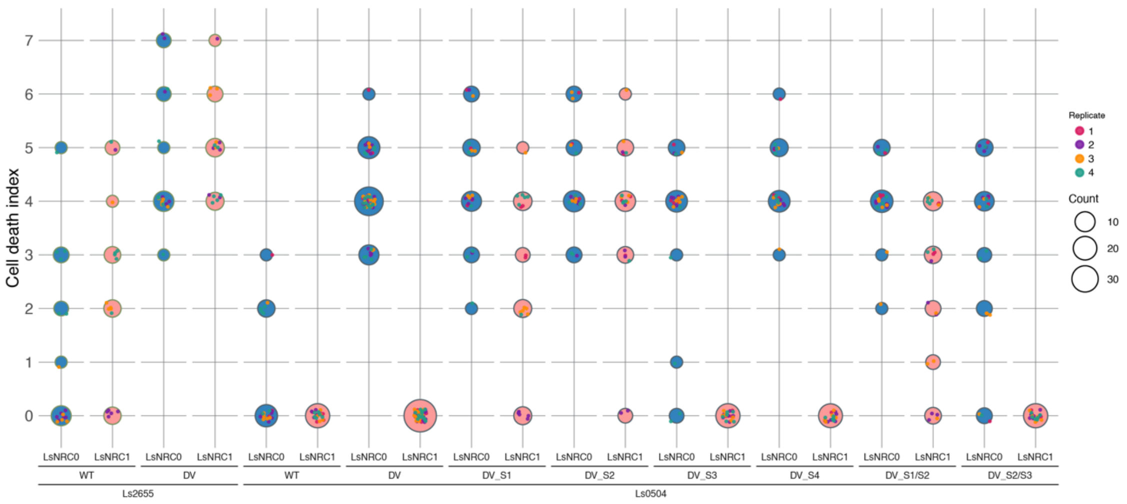


**Figure S15. Quantitative analysis of cell death assays for Ls0504 interface stretch swap chimeras.**

Quantitative analysis of cell death assays shown in **Fig. 8D**. Cell death was scored using a modified 0–7 scale at 3–5 days post agroinfiltration (*50*). Scores are shown for Ls2655 (WT and DV) and Ls0504 (WT, DV, and DV chimeras carrying individual or combined NB-domain interface stretch swaps from Ls2655: S1, S2, S3, S4, S1/S2, and S2/S3), each tested with LsNRC0 and Ast-LsNRC1. Dot size is proportional to the number of samples with the same score (Count). Colors indicate biological replicates (1–4). Data represent four biological replicates.

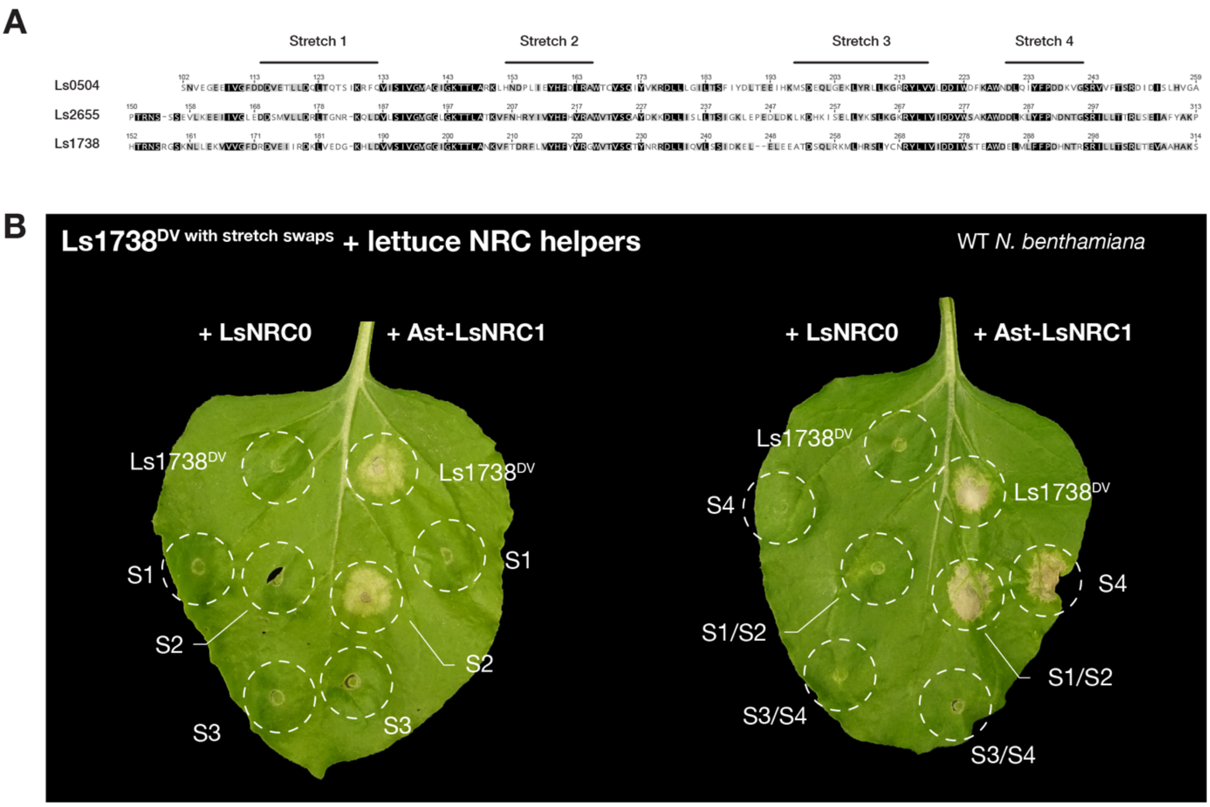

**Figure S16. Interface stretch swaps in Ls1738 do not expand helper compatibility.**

(**A**) Sequence alignment of NB-domain interface stretches (S1–S4) from the Clade 3 sensor Ls1738 and the promiscuous Clade 2 sensor Ls2655. Polymorphic residues selected for stretch swaps are indicated. (**B**) Representative WT *N. benthamiana* leaf images (brightfield) showing cell death after co-expression of LsNRC0 or Ast-LsNRC1 with Ls1738$^{DV}$ or the indicated Ls1738 $^{DV}$ chimeras carrying interface stretch swaps from Ls2655 (S1, S2, S3, S4, S1/S2, S3/S4, as indicated). Ls1738 $^{DV}$ served as a positive control for Ast-LsNRC1-dependent activation. Dashed circles indicate infiltration zones. None of the chimeras acquired the ability to signal through LsNRC0, and several swaps additionally abolished signaling through the native helper Ast-LsNRC1. The experiments were repeated three times with at least six technical replicates per repeat, with similar results in all cases.

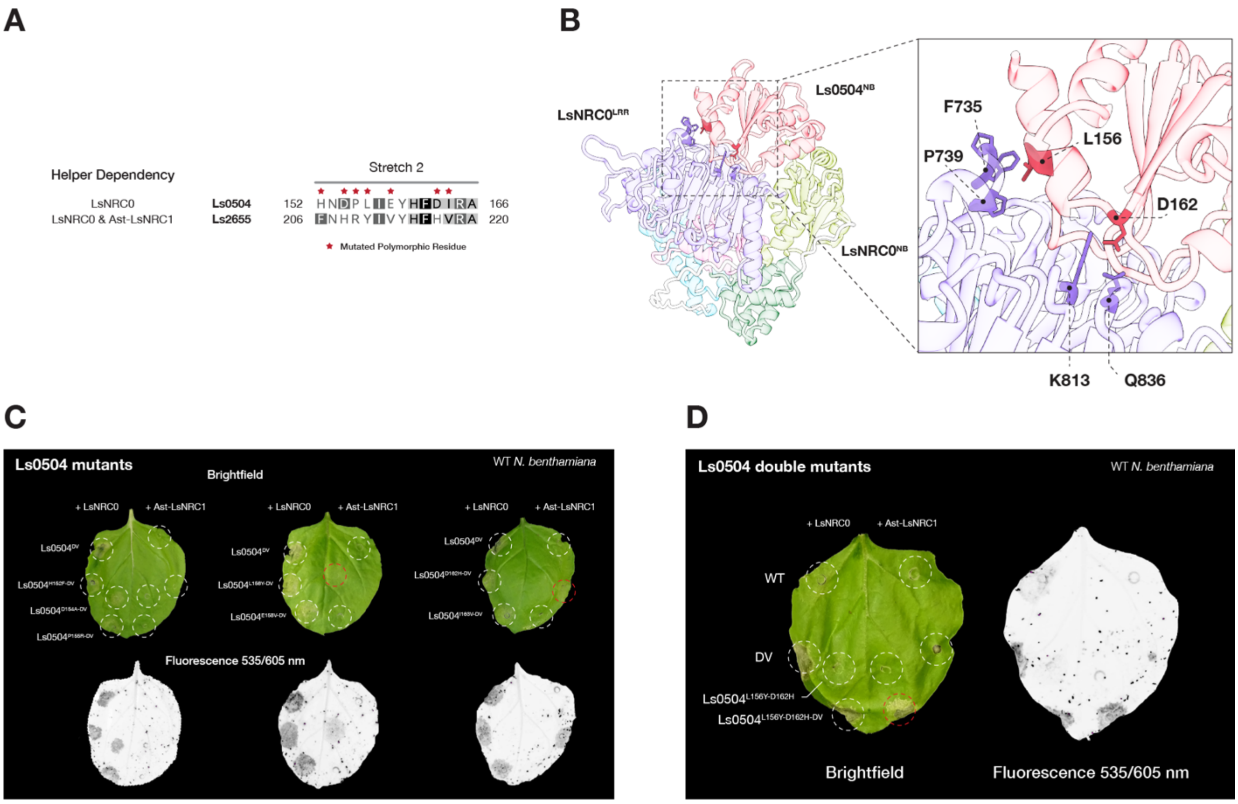


**Figure S17. Two point mutations in the Ls0504 NB domain expand helper compatibility.**

(**A**) Sequence alignment of NB-domain Stretch 2 from the Clade 1 sensor Ls0504 and the promiscuous Clade 2 sensor Ls2655, showing their respective helper dependency profiles. Polymorphic residues selected for mutagenesis are indicated with red stars. (**B**) AlphaFold 3-predicted structure of the Ls0504$^{NB}$–LsNRC0 complex highlighting the positions of L156 and D162 within the sensor–helper interface. The zoomed inset shows the predicted contacts between these two Ls0504 residues and LsNRC0 NB (F735, P739) and LRR (K813, Q836) domain residues. (**C**) Representative WT *N. benthamiana* leaf images (brightfield and fluorescence 535/605 nm) showing cell death after co-expression of LsNRC0 or Ast-LsNRC1 with Ls0504$^{DV}$ or the indicated Ls0504$^{DV}$ Stretch 2 point mutants carrying individual polymorphisms from Ls2655. Ls0504$^{DV}$ co-expressed with LsNRC0 and Ast-LsNRC1 served as positive and negative controls for helper-dependent activation, respectively. Dashed circles indicate infiltration zones; red outlines indicate gain of Ast-LsNRC1-dependent cell death. The experiments were repeated three times with at least six technical replicates per repeat, with similar results in all cases. (**D**) Representative WT *N. benthamiana* leaf images (brightfield and fluorescence 535/605 nm) showing cell death after co-expression of LsNRC0 or Ast-LsNRC1 with Ls0504 (WT), Ls0504$^{DV}$, Ls0504$^{L156Y\text{-}D162H}$, or the autoactive double mutant Ls0504$^{L156Y\text{-}D162H\text{-}DV}$. Dashed circles indicate infiltration zones; red outlines indicate gain of Ast-LsNRC1-dependent cell death. The experiments were repeated three times with at least six technical replicates per repeat, with similar results in all cases.

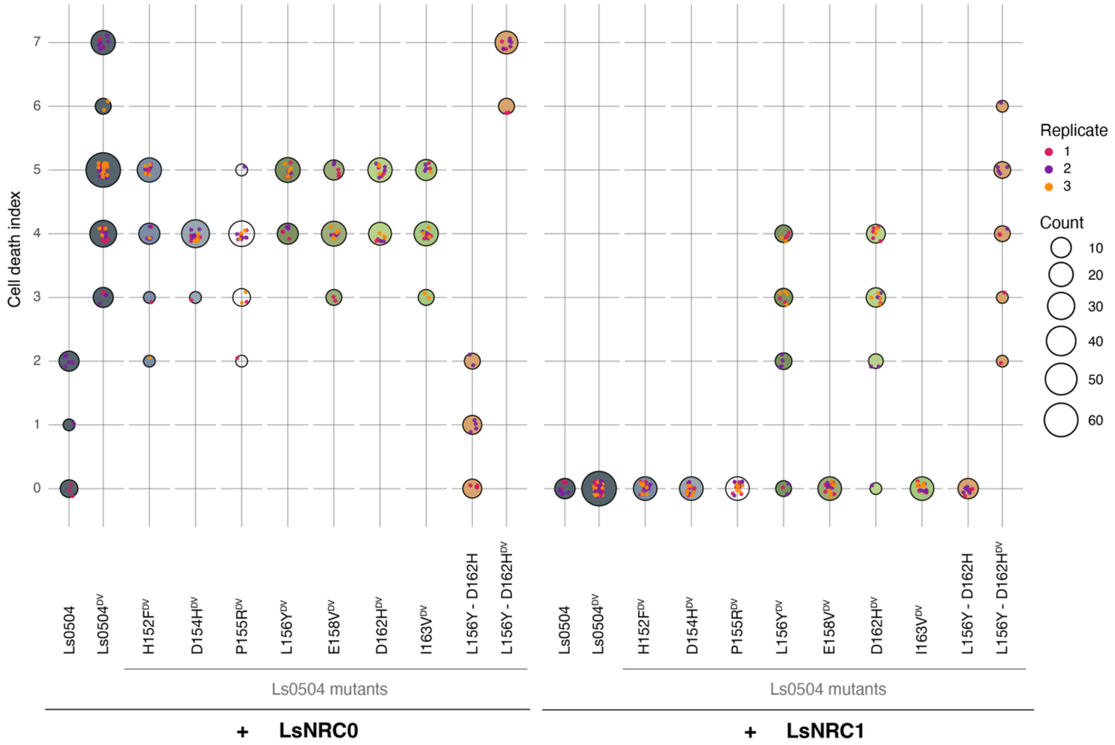


**Figure S18. Quantitative analysis of cell death assays for Ls0504 NB-domain point mutants.**

Quantitative analysis of cell death assays shown in Fig. S16C–D. Cell death was scored using a modified 0–7 scale at 3–5 days post agroinfiltration (*50*). Dot size is proportional to the number of samples with the same score (Count). The left panel shows scores for assays with LsNRC0, and the right panel shows scores for assays with Ast-LsNRC1. Data represent three biological replicates.
